# Optimal trade-off between boosted tolerance and growth fitness during adaptive evolution of yeast to ethanol shocks

**DOI:** 10.1101/2024.01.03.574056

**Authors:** Ana P. Jacobus, Stella D. Cavassana, Isabelle I. Oliveira, Joneclei A. Barreto, Ewerton Rohwedder, Jeverson Frazzon, Thalita P. Basso, Luis C. Basso, Jeferson Gross

## Abstract

The design of selection protocols to obtain bioethanol yeasts with higher alcohol tolerance poses the challenge of improving industrial strains that are already robust to high ethanol levels. Furthermore, yeasts subjected to mutagenesis and selection, or laboratory evolution, often present adaptation trade-offs wherein higher stress tolerance is attained at the expense of growth and fermentation performance. We conducted an adaptive laboratory evolution by challenging four populations (P1–P4) of the Brazilian bioethanol yeast, *Saccharomyces cerevisiae* PE-2_H4, through 68–82 cycles of 2-h ethanol shocks (19%–30% v/v) and outgrowths. Colonies isolated from the final populations (P1c–P4c) were subjected to whole-genome sequencing, revealing mutations in genes enriched for the cAMP/PKA and trehalose degradation pathways. Fitness analyses of the isolated clones P1c–P3c and reverse-engineered strains demonstrated that mutations were primarily selected for cell viability under ethanol stress, at the cost of decreased growth rates in cultures with or without ethanol. Under this selection regime for stress survival, the population P4 evolved a protective snowflake phenotype resulting from *BUD3* disruption. Despite marked adaptation trade-offs, the combination of reverse-engineered mutations *cyr1^A1474T^*/*usv1Δ* conferred 5.46% higher fitness than the parental PE-2_H4 for propagation in 8% (v/v) ethanol, with only a 1.07% fitness cost in a culture medium without alcohol. The *cyr1^A1474T^*/*usv1Δ* strain and evolved P1c displayed robust fermentations of sugarcane molasses using cell recycling and sulfuric acid treatments, mimicking Brazilian bioethanol production. These results demonstrate that some alleles selected for acute stress survival may further confer stress tolerance and optimal performance under industrial conditions.

## INTRODUCTION

Ethanol is a biofuel produced by *Saccharomyces cerevisiae* via the fermentation of hexose sugars derived from feedstocks, such as corn, sugarcane, and beet [1]. Further, this yeast species is the dominant organism driving fermentation during the manufacture of historically and economically important beverages, including wine, beer, and sake [2]. One of the reasons that *S. cerevisiae* predominates in alcoholic fermentations is related to its metabolic preference for ethanol production over respiratory processes, even in the presence of oxygen (i.e., the Crabtree effect). In addition, *S. cerevisiae* has an exceptional tolerance to life-restricting ethanol levels that occur toward the end of fermentations, potentially allowing this species to outcompete other microorganisms for ecological dominance [2].

Ethanol injures organisms primarily by disrupting the phospholipid structure of the cell membrane, which increases permeability and dissipates the electrochemical gradient [3, 4]. Damage to the cell membrane hinders the function of protein transporters and uptake of nutrients. Moreover, ethanol causes inner-cell damage by denaturing proteins and impairing mitochondrial function, provoking the formation of reactive oxygen species [3–5]. Furthermore, ethanol has mutagenic effects by interfering with the DNA replication system [6]. Despite an intrinsically high alcoholic tolerance, *S. cerevisiae* strains used in industrial processes are inevitably exposed to the detrimental effects of ethanol build-up at the final stage of fermentation, which inhibits yeast metabolism and thereby ethanol production [7, 8].

This is essential in large-scale industrial bioprocesses, such as those in the USA and Brazil, which together contribute to >80% of world’s annual ethanol production [1]. In North America, ethanol is derived from corn starch through a fermentative batch wherein ethanol titers of 13%–15% (v/v) are usually achieved [1]. In Brazil, fermentation of sugarcane juice and molasses uses cell recycling wherein yeast cells are recovered, treated with sulfuric acid to inhibit microbial contamination, and reused in repeated rounds of fermentation to reach final ethanol concentrations of 10%–12% (v/v) [1, 9–11]. Considering these bioprocesses, yeast strains with higher ethanol tolerance could more efficiently convert sugar into alcohol, improving ethanol yields while lowering distillation costs and reducing the waste footprint [7, 8, 11, 12]. For a large-scale ethanol plant operating daily with fermentation vats having capacities >one million liters, even a minimal gain in ethanol yield per fermentation batch represents a considerable increment in annual production [11].

To increase such economic gains provided by alcohol-tolerant yeasts, how to design selection screens for isolating strains that are more suitable for the bioethanol industry? For that, a crucial point such as how ethanol tolerance is defined and quantified should be considered [13, 14]. Evaluating the growth capacity of yeast cells in the presence of high ethanol concentrations is the most commonly used approach for assessing alcoholic tolerance [13–15]. Cell propagation under restrictive ethanol levels provides a practical selection method, permitting estimation of alcoholic tolerance via spot assays, maximum specific growth rates (µ_max_) on growth curves, or competition assays against reference strains. However, higher growth rate in the presence of high ethanol concentrations has been weakly correlated with a higher ethanol production capacity in fermentations [7, 12]. Furthermore, strains displaying tolerance to moderate ethanol levels (e.g., 6%–8% v/v) under laboratory conditions, sometimes do not exhibit the best performance under higher ethanol concentrations (12%–17% v/v) reached in industrial fermentations [7]. This implies the existence of various genetic factors controlling yeast tolerance to distinct ethanol levels [7, 15]. Another drawback is that several studies conducted to assess ethanol tolerance have used laboratory strains that are usually stress sensitive, making it difficult to extrapolate the results to industrial yeasts [13].

The ability to produce ethanol under very high gravity fermentations (reaching ethanol titers of 15%–17% v/v) is a more realistic proxy for assessing ethanol tolerance under industrial conditions [12, 13]. However, using high ethanol production capacity as a benchmark in selection screenings is rather unpractical because it requires fermentations lasting for several days and quantification of ethanol accumulating in several individual strains [12]. Alternatively, proliferation at moderate (e.g., 10%–12% v/v) [7] and high (17% v/v) [8] ethanol concentrations has been used for selecting strains demonstrating high ethanol accumulation. Through these various approaches, quantitative trait loci (QTLs) mapping studies have revealed the genetic basis of maximum ethanol production capacity in yeasts by identifying specific causative alleles related to genes *MKT1*, *APJ1*, *VPS70*, and *KIN3* [8, 12, 16].

Another commonly used parameter to evaluate ethanol tolerance is cell survival at prohibitive ethanol concentrations [7, 13, 14]. Ability to survive a few hours of exposure to very high ethanol levels (hereafter referred to as ethanol shocks) provides an easy method for screening tolerant yeasts [7, 17]. Selection schemes using ethanol shocks can be applied in bulk, and high lethality facilitates selective sweeps, wherein adaptive genetic variants swiftly reach dominance in a population [17, 18]. However, yeasts selected to withstand drastic treatments with 18%–22% (v/v) ethanol for 16 h have been prone to detrimental side effects and exhibited inferior fermentation and maximum ethanol accumulation capacities [7]. Circumstances wherein an organism increases fitness in a condition at the expense of a fitness reduction in another are often observed in selection screens and adaptive laboratory evolution (ALE) experiments [17–19]. Adaptation to severe stresses is even more prone to trade-off effects associated with low growth fitness; cells tend to reallocate energy resources, normally used for propagation, to overcome a life-threatening condition [20–22]. Therefore, the main challenge of genetic screenings and ALE schemes is isolating stress-resistant yeasts strains that exhibit good growth and fermentation performance [7, 17, 19, 23].

Herein, we used an ALE protocol that combined ethanol shocks with a growth recovery period to balance the selection of strains able to tolerate acute ethanol treatments with minimal losses in growth fitness. ALE is used to isolate alcohol-tolerant yeasts, mostly via propagation in the presence of ethanol [15, 24–33]. Although ALE experimental designs also used drastic ethanol treatments [34, 35], numerous questions regarding the outcome and utility of this selection method remain unanswered. These questions pertain to the extent to which attained tolerance to ethanol shocks causes a decline in growth fitness. Furthermore, much information regarding the mutational spectrum underlying tolerance to drastic ethanol treatments remains unknown. Do all the adaptive alleles for survival in ethanol shocks negatively impact growth rates, or do some mutations benefit growth in the presence of ethanol, supporting higher ethanol yields during fermentation? An important point to interrogate is whether yeast selection via harsh ethanol treatments could confer robust strains with superior performance in bioprocesses involving various stress factors; this question is pertinent to the Brazilian bioethanol production from sugarcane substrates, which involves high cell densities wherein yeasts are exposed to fluctuations in osmotic pressure, temperature, microbial contamination, toxic compounds, and a sharp sulfuric acid shock intercalating sequential fermentation cycles [1, 10, 36].

## RESULTS

### Yeast ALE with drastic ethanol treatments

We founded four haploid ALE populations (P1–P4, Fig 1A) of the Brazilian bioethanol strain *S. cerevisiae* PE-2 [36]. P1, P2, and P3 are *MATa* populations derived from the strain PE-2_H4 (S1 Table), whereas P4 originated from an *MATα* spore resulting from a cross between PE-2_H3 and PE-2_H4 [9]. For ALE, P1–P4 were subjected to several cycles consisting of ethanol shocks for 2-hours followed by recovery periods of 2–4 days growth in liquid yeast extract-peptone plus 2% sucrose (YPS) (Fig 1A). To challenge the yeast adaptation, the 2-hour shocks entailed progressive increments of ethanol concentration from 19% up to 30% (v/v) (Fig 1B). At the end of evolution experiment, a single colony (clone) was selected from solid YPS medium cultures (plus 8% v/v ethanol) of each population (Fig 1A). Isolated clones P1c–P4c were subjected to whole-genome sequencing (WGS), and P1c–P3c were phenotypically characterized. The isolated P4c was excluded from most fitness analyses as the P4 population evolved in a snowflake-type aggregation (see below), which precluded the detection of individual cells through flow cytometry and colony-forming assays.

**Fig 1.**
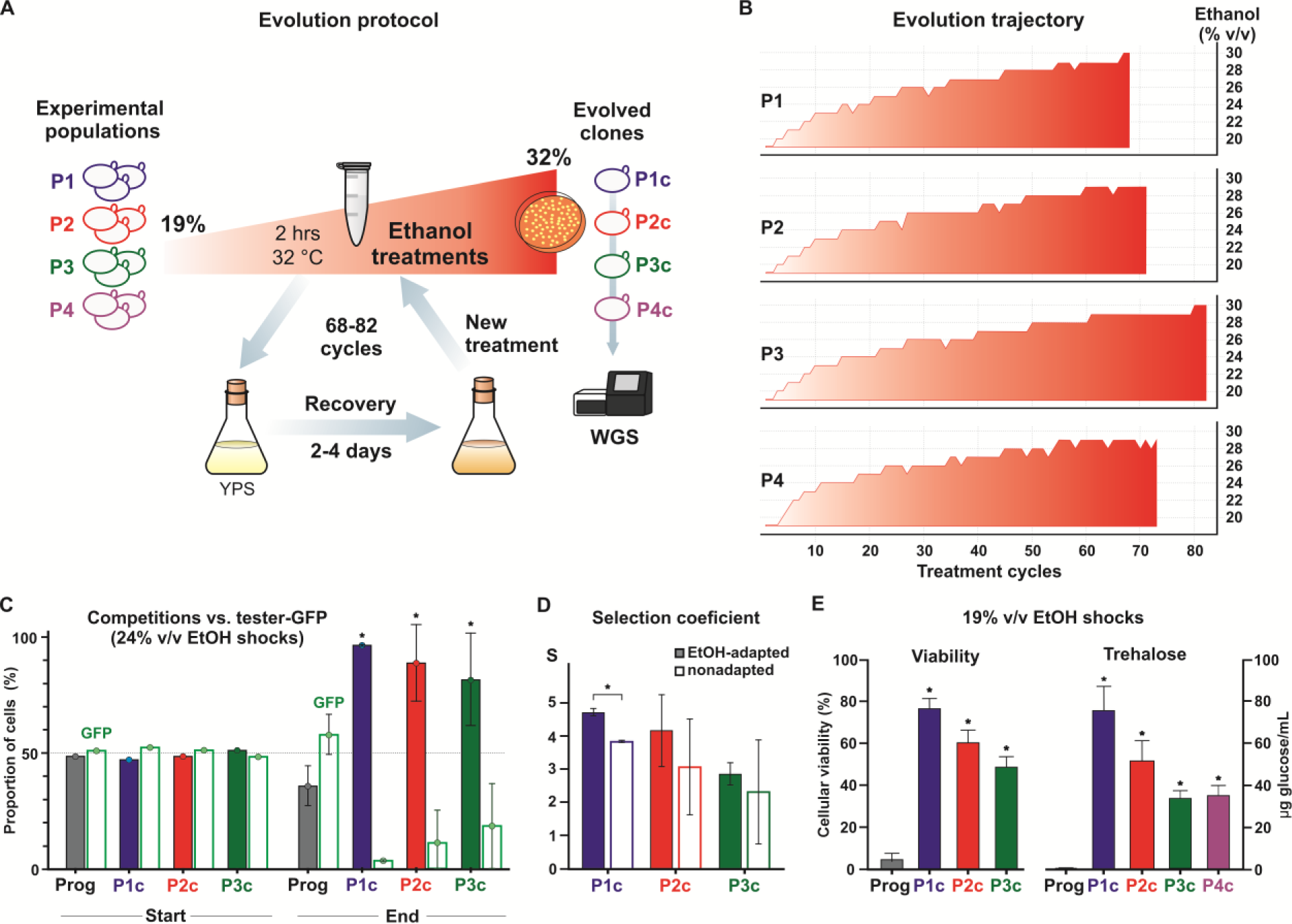
Adaptive evolution experiment (ALE) with ethanol treatments. (**A**) The scheme of ALE wherein four yeast populations (P1–P4) were subjected to repeated cycles of 2-hour ethanol shocks (about 10^8^ cells resuspended in 1 mL PBS 1X buffer with ethanol) and outgrowth in yeast extract-peptone plus 2% sucrose (YPS). At the end of the experiment, one colony in solid medium with 8% (v/v) ethanol was isolated from each population for whole-genome sequencing (WGS). (**B**) The increase in ethanol concentrations (red) is plotted according to the number of shock/recovery cycles for each population. (**C**) The progenitor (Prog.), P1c, P2c, and P3c were first propagated in YPS for five cycles, then they were separately mixed in equal proportions with a reference strain expressing a green fluorescence protein (tester-GFP). A competition assay was performed through one cycle of ethanol shock (24% v/v for 2 hours) and growth recovery in YPS. Flow cytometry recorded the proportion of cells for each competitor just before the ethanol shock (Start) and at the stationary phase following recovery growth (End). (**D**) Selection coefficients (S) were calculated for nonadapted competitors (panel **C**, S2 Table) and for cells adapted by five progressive ethanol treatments (EtOH-adapted, see text and methods). The S > 0 indicates that the cell numbers of evolved clones largely exceeded those of the tester-GFP at the competition end point. S was normalized with data obtained for the progenitor (see methods, S2 Table). (**E**) For cellular viability estimation, after 19% (v/v) ethanol shocks, cells were diluted and plated into solid medium for counting the resulting colonies. For trehalose quantification, postshocked cells were recovered by growth in YPS. Quantification was obtained from 10^8^ cells by the enzymatic conversion of trehalose into glucose (see methods). (*) p<0.001, one way ANOVA followed by Bonferroni post-test for multiple comparisons.

To test whether the increase in ethanol tolerance observed during ALE (Fig 1B) was a stable trait, we propagated P1c, P2c, and P3c over five passages in YPS without ethanol. Then, competition assays were performed wherein the progenitor and evolved clones P1c–P3c were separately mixed in equal proportions with a green fluorescent protein (GFP)-expressing PE-2_H4 strain (tester-GFP). The competitors were exposed to a 24% (v/v) ethanol shock for 2 hours, followed by a recovery in YPS until the stationary phase. The proportion of each competitor at initial and final points of the assay was determined using a flow cytometer (Attune NxT, Thermo Fisher Scientific) (Fig 1C and S2 Table). The ALE progenitor showed a slight decrease in proportion when competing with the tester-GFP, whereas P1c–P3c outcompeted the tester-GFP by a large margin (Fig 1C). The selection coefficients of the evolved clones demonstrated a more pronounced improvement in fitness than the progenitor (Fig 1D, S2 Table). We repeated the fitness assays by subjecting all competitors to a preacclimation (i.e., progressive exposures to 15%, 15%, 20%, 20%, and 22% v/v ethanol) before the 24% (v/v) ethanol shock. After ethanol treatment and recovery, the selection coefficients for preadapted P1c–P3c were only slightly higher than those of the nonadapted clones, and only significant for P1c (Fig 1D). Ethanol tolerance of the evolved clones was also characterized by a marked increase in survival rates following 2-h treatments with 19% (v/v) ethanol (Fig 1E). Interestingly, following exposure to ethanol, trehalose accumulation by P1c–P4c was proportional to their postshock fitness and survival rates, indicating a common mechanism for ethanol tolerance (Fig 1E). For example, trehalose content was highest in P1c, which also displayed the highest survival rate and selection coefficient following the ethanol shock.

### WGS of evolved clones

Illumina MiSeq platform was used for sequencing the genomic DNA of P1c, P2c, P3c, and P4c. Variant calling for P1c, P2c, and P3c was accomplished using the PE- 2_H4 genome as a reference, whereas the P4c read mappings to both PE-2_H3 and PE-2_H4 parental genomes [9] were used to identify de novo single nucleotide polymorphisms (SNPs) not present in either reference genomes. We cataloged 46 mutations in total (Fig 2A and S3 Table). We identified potentially adaptive alleles related to genes that were mutated in the various populations (Fig 2A–2C). The gene *ATH1* (encoding the acid trehalase) had defective mutations in P1c, P2c, and P3c (Fig 2A–2C). *CYR1* (P1c and P2c), *PTR2* (P3c and P4c), *MDS3*, *ROM2*, and *USV1* (P1c and P3c) were the other genes with mutations appearing in two different clones. Various mutated genes acting on the same metabolic or signaling pathway imply parallelism. Mutations affecting genes related to the cAMP/PKA pathway (i.e., *CYR1*, *MDS3*, *PMD1*, *RAS2*, and *IRA2*) are dominant in our dataset. In addition to the three inactive *ath1* alleles, *NTH1* (encoding the neutral trehalase) had a frameshift mutation in P3c, indicating that blocked trehalose catabolism was under selection during ALE. This criterial triage of mutations was important for guiding our reverse-engineering program described below.

**Fig 2.**
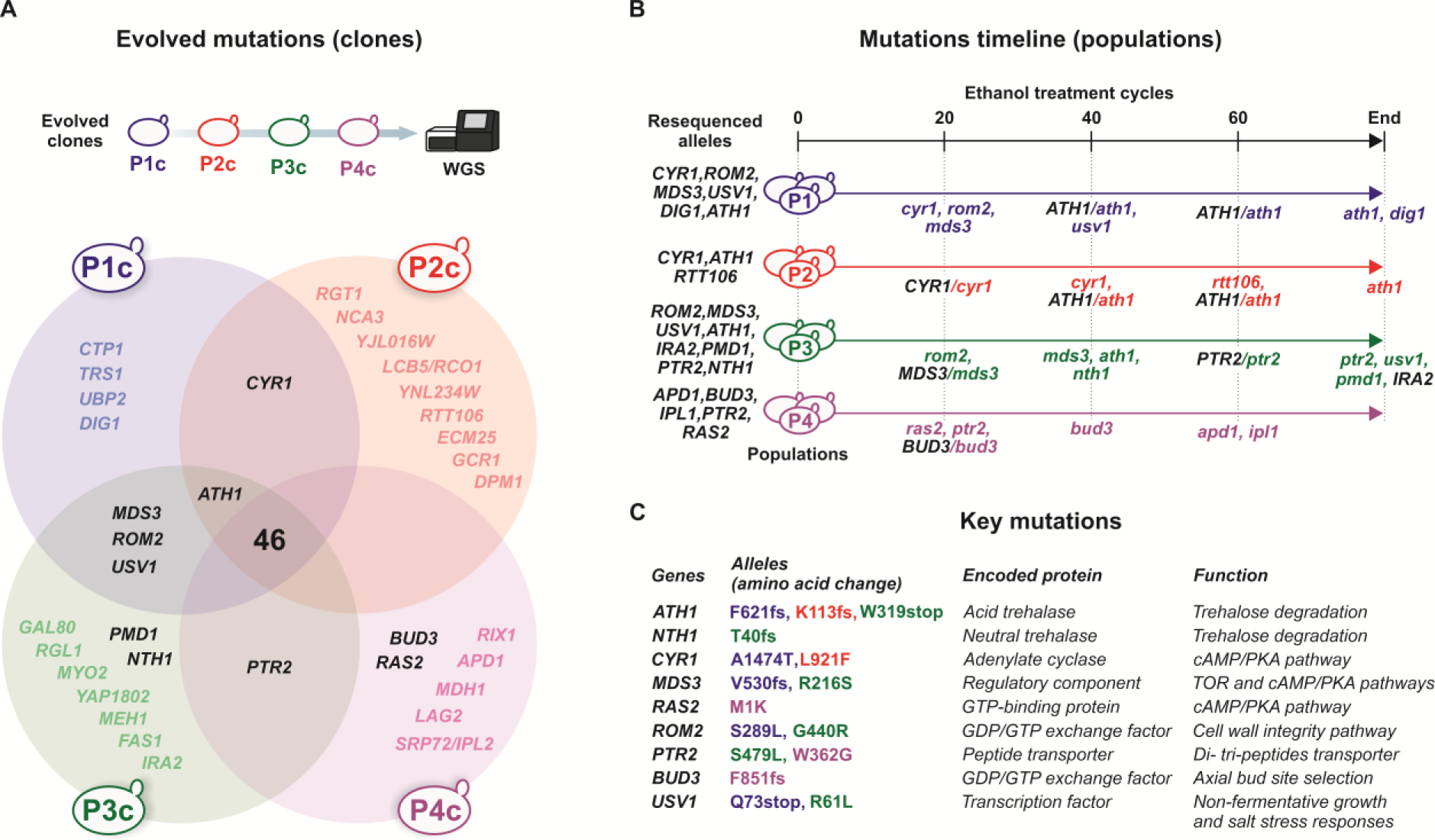
WGS data from P1c, P2c, P3c, and P4c. (**A**) Venn diagram displaying unique mutated genes found for each clone, and genes that had mutations in more than one clone (intersections between circles). Highlighted in bold back fonts are genes whose mutated alleles are important for this study. (**B**) Sanger sequencing of selected alleles in progenitors, final, and intermediate populations. Gene names in uppercase indicate the alleles found in progenitors. Gene names in lowercase indicate the mutated alleles at the point they were first detected by Sanger sequencing. When progenitor and evolved alleles are depicted, the evolved alleles are at an intermediate frequency on that sampled point. (**C**) List of key mutated genes and their encoded proteins important for further analysis in this work.

We confirmed a selected set of mutations via Sanger sequencing of PCR fragments amplified from the progenitor and final populations (Fig 2B and S1 Appendix). With the exception of the *ira2* allele (observed in the P3c, but not in the P3 final population), we confirmed that all mutations were present in the corresponding final populations, whereas, as expected, progenitors had wild-type alleles. Furthermore, we Sanger sequenced the same loci in intermediate ALE populations sampled following ethanol shocks 20, 40, and 60 (Fig 2B and S1 Appendix), allowing us to identify when the mutation became dominant in the population (i.e., there was a unique peak in the corresponding position on the chromatogram) or when alleles had intermediate frequencies in the population (i.e., overlapping peaks corresponding to the wild-type and evolved alleles were present on the chromatogram). Generally, in ALE experiments, mutations related to large fitness effects appear early during ALE experiments [17, 18]. Therefore, in our evolution experiment, *cyr1*, *mds3*, *rom2*, *ras2*, and *ptr2* alleles (all present at shock 20) are the most likely drivers of ethanol tolerance (Fig 2B). Interestingly, although three *ath1* defective alleles appeared in different populations, they emerged only later during ALE (Fig 2B), suggesting that the earlier mutations (e.g., *cyr1*, *mds3*, and *pmd1* that affected the cAMP/PKA pathway and increased trehalose accumulation) could have potentiated their appearance.

### Evolved clones increased Msn2/4-mediated stress responses

Mutations affecting the cAMP/PKA pathway occurred in all four ALE populations. Disruptive mutations, such as those present in *RAS2* (P4), *MDS3* (P2 and P3), *PMD1* (P3), and nonsynonymous mutations in *CYR1* (P1 and P2), can downregulate protein kinase A (PKA) and its downstream responses (Fig 3A) [37–40]. If PKA is inhibited, transcription factors Msn2/4 are retained in an unphosphorylated form and shuttle into the nucleus to drive the expression of stress-responsive genes [40]. One such stress-regulated gene is *HSP12*, which encodes a heat-shock protein, whose expression when fused with GFP has been used as a standard biosensor to measure Msn2/4 regulation (Fig 3A) [37]. We introduced a chromosome-integrated *HSP12-GFP* biosensor into P1c, P2c, P3c, and in the parental PE-2_H4 backgrounds. Furthermore, the biosensor was introduced into a strain reverse-engineered with the P1c allele *cyr1^A1474T^* (Fig 3B-3D). Relative to the PE-2_H4 reference, the P1c–P3c displayed a constitutive upregulation of the *HSP12-GFP* expression; the expression was more pronounced in the P2c (Figs 3B and 3C, S1 Fig). Moreover, the *cyr1^A1474T^* reverse-engineered strain exhibited a higher *HSP12-GFP* expression than that of its parent PE-2_H4. The higher GFP expression in evolved clones than in the parental strain seems to be constitutive, and not dependent on the ethanol treatment (Fig 3D), supporting the hypothesis that evolved clones, carrying mutations in *CYR1*, *RAS2*, *MDS3*, and *PMD1* (Fig 2), display a low PKA phenotype wherein Msn2/4p is activated and upregulates the expression of stress-responsive genes [37–41]. The boost in trehalose biosynthesis, demonstrated by the higher content of trehalose observed in the evolved P1c–P4c (Fig 1E), is typical of strains with low PKA activity.

**Fig 3.**
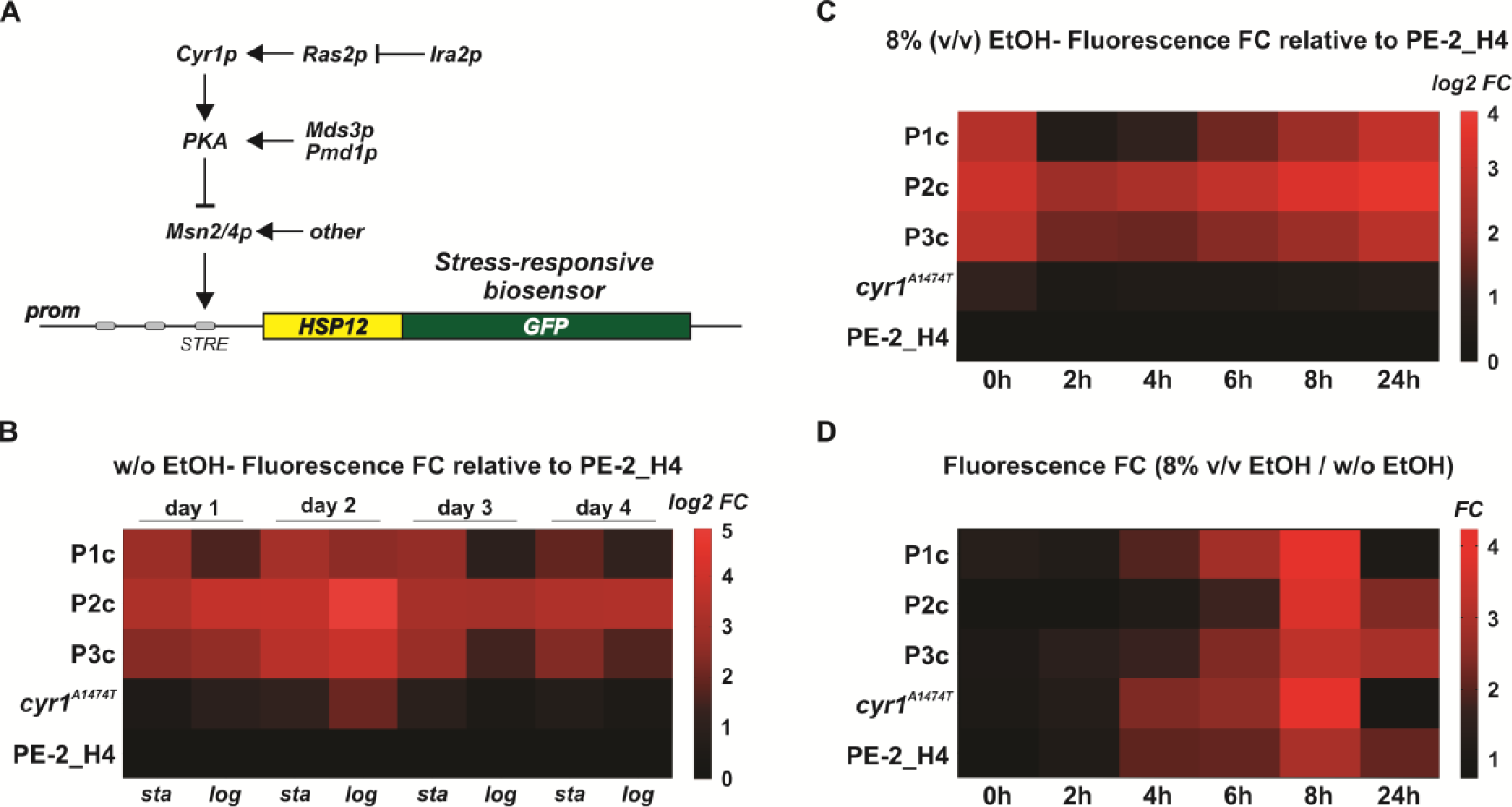
Upregulation of Msn2/4p-mediated stress responses in evolved clones. (**A**) Scheme depicting the *HSP12-GFP* biosensor to measure Msn2/4p-mediated stress responses. Msn2/4p binds to STRE elements in the *HSP12* promoter, inducing biosensor expression. The GFP fluorescence signal is proportional to the Msn2/4p activation. Components of the cAMP/PKA pathway controlling the Msn2/4p function have been shown. Mutations, found through WGS of evolved clones, affect molecular components controlling the PKA, putatively resulting in upregulation of Msn2/4p-mediated transcription. The biosensor was integrated into the P1c, P2c, P3c, and *cyr^1A1474T^* backgrounds. (**B**) Time course of GFP fluorescence signal during four serial transfers in YPS (without ethanol). Fluorescence measurements were obtained at the stationary (*sta*) and logarithmic (*log*) growth phases. P1c, P2c, P3c, and *cyr^1A1474T^* data are expressed as log_2_ of fluorescence fold changes (FC) relative to the PE-2_H4 signal at the same time point. (**C**) Time course of GFP fluorescence along 24 hours in cells propagating in 8% (v/v) ethanol. Data represent log_2_ FC relative to the PE-2_H4 fluorescence at the same time point. (**D**) Induction of GFP expression at 8% (v/v) ethanol relative to cells propagating in the absence of ethanol. Relative fluorescence FC at 8% (v/v) ethanol were obtained by comparing with the cells propagating without ethanol at the same time point. Values indicate the mean of measurements obtained from three independent replicates.

### Gains in tolerance to ethanol shocks usually involve growth fitness losses

We reverse-engineered key evolved mutations into the parental background PE- 2_H4 (S1 Text, S1 and S4 Tables). Loss-of-function mutations (i.e., premature stop codons or frameshift mutations) were mimicked by insertional disruption of genes with the *MX* cassette [42]. In other instances, coding frame deletions were edited using the CRISPR/Cas9 EasyGuide method [43]. Furthermore, point mutations were introduced via CRISPR/Cas9. Thus, series of single- and double-mutants was constructed and tested via three different competition assays against the tester-GFP (Fig 4A) (S5 Table).

**Fig 4.**
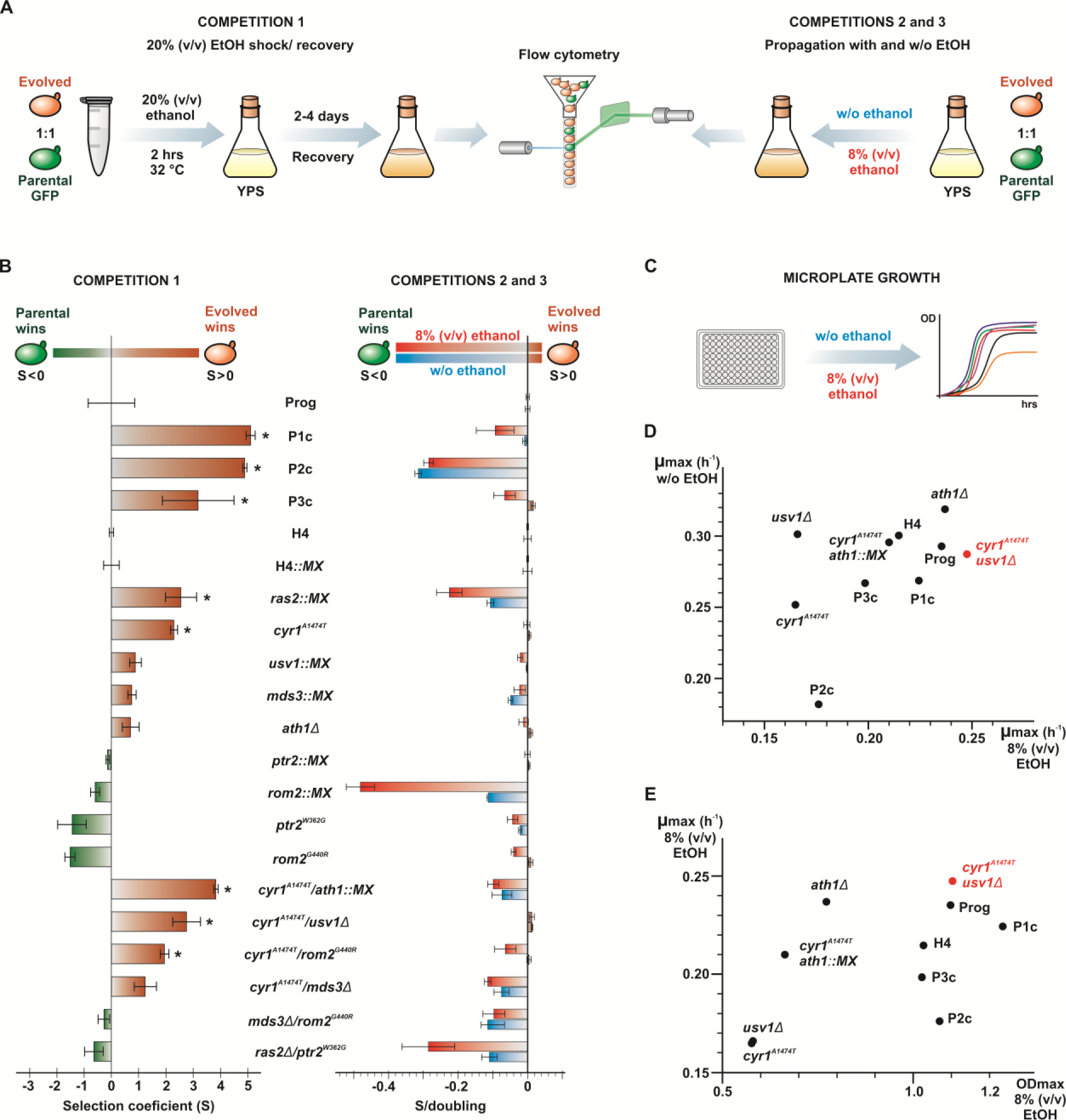
Screenings for ethanol tolerance with ALE clones and reverse-engineered strains. (**A**) Scheme showing three assays wherein a strain under test was competed against a GFP-marked parental PE-2_H4 (tester-GFP). Competition 1 that mimics our ALE approach based on 20% (v/v) ethanol treatments for 2 hours is followed by recovery growth and flow cytometry measurements of fluorescence-tagged and nonfluorescent cells. Competitions 2 and 3 were based on a single cycle of propagation in liquid YPS medium without ethanol (approximately 8–10 doublings) and with 8% (v/v) ethanol added to the YPS culture medium (approximately 5–7 doublings), respectively. (**B**) Initial and final ratios of competitors (measured by flow cytometry, S5 Table) were used to calculate the selection coefficients (S) of each tested strain for assays 1 (left), 2, and 3 (right). Numbers were obtained from three replicates. Values of S were normalized with values obtained for competitions of the corresponding parental strains against the tester-GFP. For competitions 2 and 3 the S values are expressed per cell doubling. A positive S indicates that the strain under test has a higher fitness than its parental. Statistical significance of S values obtained for mutant strains was assessed in comparison to their respective parentals. (*) p<0.05, one way ANOVA followed by Bonferroni post-test for multiple comparisons. (**C**) A selected group of strains was also evaluated through microplate growth assays in 8% (v/v) ethanol and normal YPS medium. (**D**) Maximum specific growth rates (µmax) obtained in 8% ethanol and without ethanol were plotted. The strain *cyr1^A1474T^/usv1Δ* is highlighted (red) for having the best growth performance in 8% (v/v) ethanol. (**E**) Plotting of µ_max_ vs. maximum optical density (OD_max_) at the plateau of growth curves in 8% (v/v) ethanol. Strain *cyr1^A1474T^/usv1Δ* (red) shows optimal combination of µmax and OD_max_.

For competition assay 1, we used a cycle of 20% (v/v) ethanol shock for 2 h, followed by a recovery growth in YPS (Fig 4A and 4B). Under this condition mimicking our ALE protocol, evolved P1c, P2c, and P3c as well as individual mutants *cyr1^A1474T^*, *mds3::MX*, and *ras2::MX*, related to the cAMP/PKA pathway, exhibited higher fitness (i.e., positive selection coefficient, S > 0) than their respective parentals (Fig 4B). The *ath1Δ* deletion and *usv::MX* disruption were also adaptive. A combination of *cyr1^A1474T^* plus *ath1::MX and cyr1^A1474T^*/*usv1Δ* displayed synergistic effects. However, no positive fitness contributions were observed for *rom2^G440R^* and *ptr2^W362G^* even when they were combined with other alleles. These observations were contrary to our expectations, as *ROM2* and *PTR2* displayed mutations that emerged early during ALE in two different populations (Fig 2A and 2B).

In addition to testing yeast tolerance to a transient 2-hour pulse of high alcohol concentration (20% v/v), we probed adaptation for propagation in constant ethanol exposure. Competition assay 2 involved inoculation of competitors in YPS at an 8% (v/v) ethanol concentration and propagation for up to 3 days until the stationary phase (Fig 4A and 4B). Furthermore, we analyzed the growth for 24-hours wherein competitors were inoculated in YPS without ethanol (competition assay 3). Most evolved clones and reverse-engineered strains had a notable fitness loss when propagated in the presence of 8% (v/v) ethanol and also without ethanol (Fig 4B). Despite being highly adapted to ethanol shocks, the evolved clones P1c–P3c and mutants *cyr1^A1474T^*/*ath1::MX*, *cyr1^A1474T^*/*mds3Δ*, *cyr1^A1474T^*/*rom2^G440R^*, *ras2::MX*, and *mds3::MX* exhibited a pronounced decline in fitness when propagated under 8% (v/v) ethanol, and most of them also had a decreased performance when grown in normal YPS. These observations indicate trade-off effects for mutations that cause adaptation to transient and drastic ethanol treatments; i.e., although mutations promote cell survival in a punishing environment, they imposed a fitness cost for propagation under mild or nonstressful conditions.

However, *cyr1^A1474T^*, *usv1::MX*, *ath1Δ*, and *cyr1^A1474T^*/*usv1Δ* maintained growth fitness similar to that of their parental strains, indicating that an equilibrium between acute stress tolerance and propagation fitness might be attainable. To gain further insights into the tested strains, we performed microplate growth assays involving the PE-2_H4 parental, the evolved clones P1c–P3c, and the best-performing reverse-engineered strains that could balance ethanol shock tolerance with cell-doubling fitness (Fig 4C and 4D, S6 Table). The *ath1Δ* mutant displayed excellent growth rates in YPS with and without ethanol; however, it had a very low maximal optimal density (OD_max_) in the presence of ethanol. In contrast, the evolved clone P1c demonstrated superior biomass yield at 8% (v/v) ethanol, a selective trait that was presumably advantageous during the ALE postshock recovery growth until the stationary phase. We recorded an optimal performance for strain *cyr1^A1474T^*/*usv1Δ*, which presented the highest specific growth rate (µ_max_) at 8% (v/v) ethanol; in normal YPS it maintained a µ_max_ slightly below that observed for the parental PE-2_H4. These results indicated that our screening with ethanol shocks could select alleles that, in combination, improved yeast growth in the presence of ethanol.

### Population P4 evolved a snowflake phenotype driven by *BUD3* disruption

Yeast populations that form multicell aggregations displaying a “snowflake” appearance result from mutations that preclude the correct separation of mother-daughter cells following mitotic division [44, 45]. Generally, these mutations are related to the RAM network that involves the Ace2p transcription factor [46, 47]. Cell-to-cell attachments nucleating large clusters of yeasts often emerge in laboratory evolution experiments and in industrial settings from selections for increased sedimentation rates [44–46], or as adaptations to withstand various stresses [47]. We observed that the P4 population displayed a snowflake-type phenotype that became apparent by ethanol shock/recovery cycle number 40 (Fig 5A). This coincided with the dominance of the *bud3^F851fs^* loss-of-function allele caused by a single nucleotide deletion (Fig 5A and 5B, S2 Table). Although this mutation was present at low frequency at sampled point 20, it prevailed following ethanol treatment 40 (Fig 5B, S1 Appendix). Bud3p acts as a determinant for axial bud site selection and localizes to the bud neck contractile ring during mitosis [48, 49]. We observed that *bud3* insertional knockout was sufficient to generate an aggregative phenotype resembling that observed for P4 (Fig 5C). Interestingly, the diploid *bud3::MX* knockout did not display the snowflake phenotype (S2 Fig); this was in accordance with the role played by Bud3p only in haploid cells wherein the component helps to set a landmark for an axial budding pattern, while diploid cells undergo a different bipolar budding program [48, 49]. Clusters of cells in P4 may represent a collective adaptation to ethanol stress; i.e., yeasts inside the cluster may be more protected from the damaging effects of alcohol, which predominantly affect the cell plasma membrane [50, 51].

**Fig 5.**
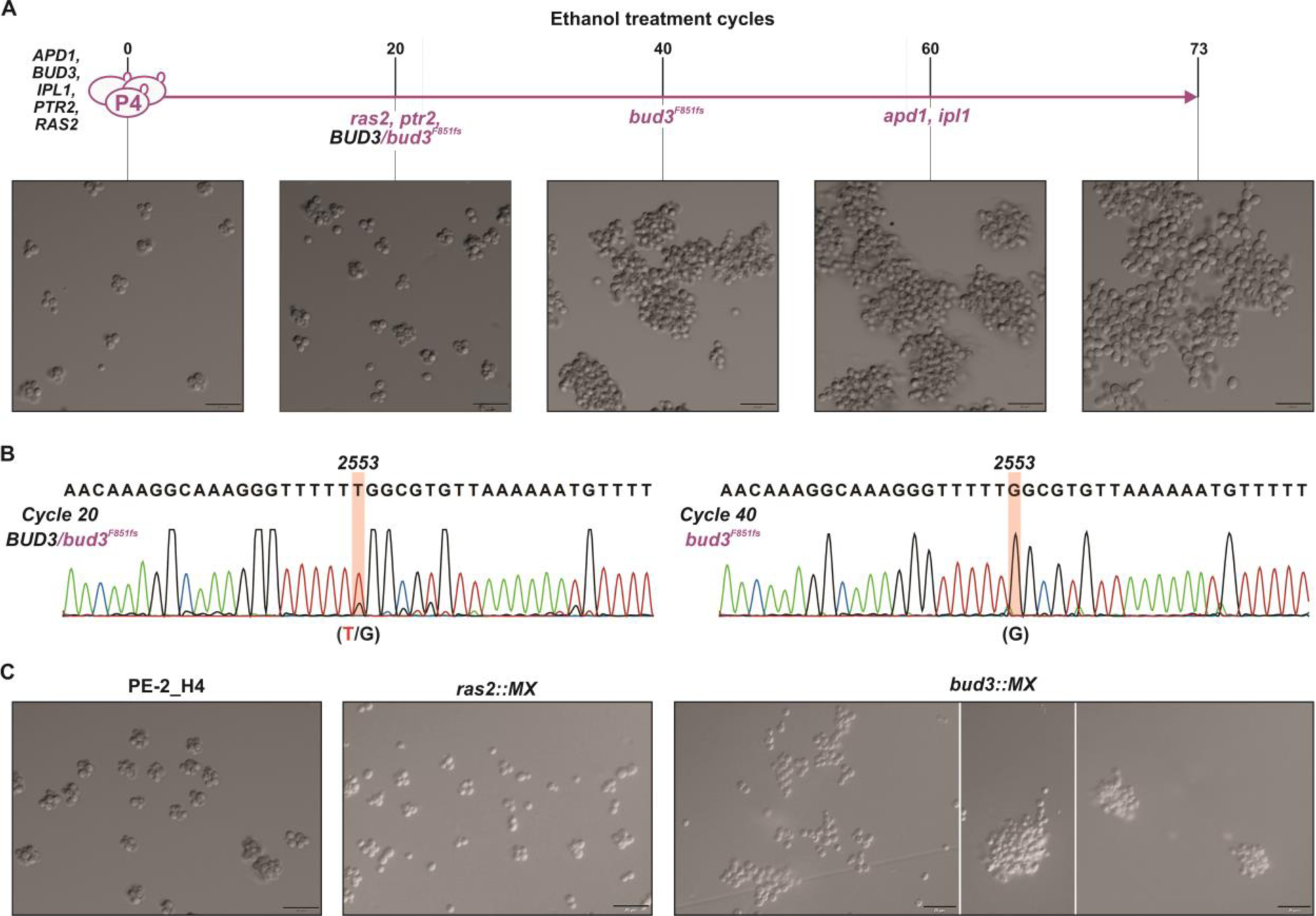
Snowflake phenotype related to P4. (**A**) Time course of ethanol shocks with population P4 and dominance of the snowflake phenotype at sampled point 40. (**B**) Although the mutation 2553DelT (*bud3^F851fs^*) was already present at sampled point 20, it became dominant by ethanol shock/recovery number 40. (**C**) Genetic underpinnings of the snowflake phenotype. The loss-of-function mutations *ras2::MX* and *bud2::MX* were constructed in the PE-2_H4 background. The cell-aggregation phenotype is associated with the insertional mutant *bud3::MX*, but not with *ras2::MX*. Magnification, ×40.

### Evolved clone P1c and the double-mutant *cyr1^A1474T^*/*usv1Δ* display optimal balance between ethanol tolerance and fermentation performance

Our screening involving transient ethanol shocks selected strains that were tolerant to an acute stress condition. The Brazilian bioethanol production system, from sugarcane juices and molasses, exposes yeasts to a dynamic fluctuation of stresses during fermentations in industrial vats [1, 10, 36]. These stresses include osmotic pressure, heat, toxic compounds, and build-up of ethanol concentration at the end of the process. Fermentation with cell recycling, used in the Brazilian ethanol production, adds a particular stressful factor; following each fermentation cycle, yeast cells are centrifuged and treated with sulfuric acid (pH = 2.5) for about 1h to kill bacterial contaminants [1, 10, 36]. This drastic acid shock constitutes a pulse of stress that circumstantially resembles the ethanol treatments used in our ALE protocol. Therefore, we explored whether tolerant yeasts selected using our shock/recovery regime exhibited a good performance in benchtop fermentations that simulate Brazilian ethanol production using cell recycling and sulfuric acid treatments [10, 36]. During fermentations using cell recycling a biomass increase of about 10% can occur. Furthermore, cell death may account for fluctuations in cell numbers from one cycle to the next [10, 11, 36, 52]. Thus, we tested the possibility of conducting competitions of strains against the reference during fermentations of sugarcane molasses using cell recycling. This would allow us to calculate cumulative selection coefficients (S), relative to the progenitor, which express the propensity of the strain to increase (or not) a viable biomass during the stressful fermentation. To test the fitness of evolved clones P1c and P3c, probed strains and the parental ALE progenitor (Prog.) were separately mixed with the PE-2_H4 tester-GFP to initiate eight cycles of sugarcane molasses fermentations (Fig 6A). During the process, cells were sampled after each cycle and the proportions of GFP-labeled to nonlabeled cells were measured using an Attune NxT flow cytometer. While P3c was outcompeted by the tester-GFP during the fermentation cycles, the increasing cumulative fitness of P1c showed that the strain was better adapted than its progenitor to the harsh conditions of ethanol fermentation with cell recycling (Fig 6A).

**Fig 6.**
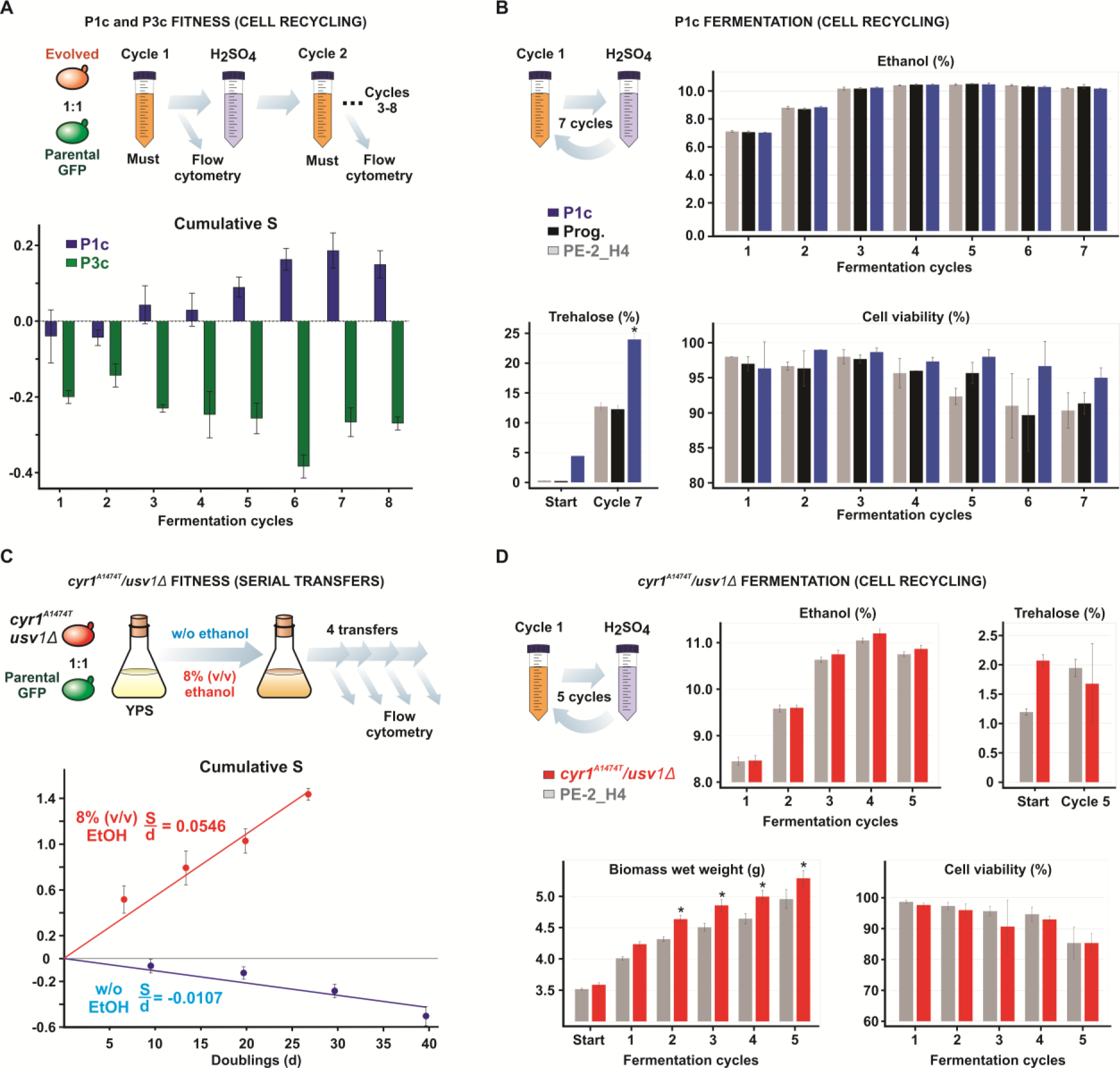
Fitness and fermentation performance of ethanol tolerant strains. (**A**) Fitness of P1c and P3c relative to the progenitor during sugarcane molasses fermentation with cell recycling. P1c, P3c, and the ALE progenitor (Prog.) were individually mixed with the tester-GFP and subjected to eight cycles of fermentations. The input of total reducing sugars (TRS) for cycles 1–8 were, respectively, 14.9%, 16.7%, 18.8%, 21.3%, 21.6%, 17.2%, 20.7%, and 20.9%. The conditions resembled the Brazilian ethanol production wherein cells are recovered by centrifugation following each fermentation and treated with sulfuric acid (H_2_SO_4_) 0.5 M for 1 hour (pH 2.5). Flow cytometry measurements allowed estimation of cumulative selection coefficient (S) after each cycle. Values of S for evolved P1c and P3c are expressed as the difference from the cumulative fitness recorded for the progenitor in a competition against the tester-GFP. (**B**) Fermentation of sugarcane molasses during seven cycles intercalated with sulfuric acid treatments. The input of TRS for cycles 1–7 were 14.9%, 18.5%, 21.4%, 22.0%, 22.0%, 21.7%, and 21.5%, respectively. Final ethanol accumulation and cell viabilities for P1c, the ALE progenitor (Prog.), and parental PE-2_H4 were quantified at each cycle. Trehalose content was estimated at the beginning and end of the experiment and was expressed as the wet cell weight percentage for each strain. (**C**) Competition assays between the *cyr1^A1474T^*/*usv1Δ* double-mutant and the tester-GFP through sequential passages in YPS with 8% (v/v) ethanol and without alcohol. Numbers are relative to the fitness calculated under the same conditions for the PE-2_H4 parental strain. Plotted are the cumulative S per number of calculated generations at the end of each passage. Selective coefficients per cell doublings (S/d) were calculated from the trendline fitted into the data points. (**D**) Fermentation of sugarcane molasses with the *cyr1^A1474T^*/*usv1Δ* double-mutant and PE-2_H4. Fermentation cycles 1–5 were performed with initial TRS content of 17.8%, 20.3%, 22.3%, 22.8%, and 22.5%, respectively. Sulfuric acid treatments were performed after each cycle. The content of ethanol, biomass, and cellular viabilities were estimated at the end of each fermentation cycle. Trehalose was quantified at the beginning and end of the fermentation. (*) p<0.05, one way ANOVA followed by Bonferroni post-test for multiple comparisons.

A separate fermentation assay conducted using individual strains demonstrated that P1c achieved ethanol titers and biomass formation similar to those of the ALE progenitor and parental PE-2_H4 (Figs 6B and S4). More importantly, P1c displayed higher cell viability after each cycle (Fig 6B), indicating that this strain was better adapted to the fermentation conditions. Trehalose accumulation, a hallmark of stress tolerance in Brazilian bioethanol yeasts [10, 36], which was already higher in the P1c compared with the parental strains at the beginning of the experiment, reached over 23% of cell wet mass at the end, nearly double the trehalose content reached by the parental strains (Fig 6B). Despite the superior traits exhibited by P1c, strains reverse-engineered with P1c mutations, *cyr1^A1474T^*, *ath1Δ*, and *cyr1^A1474T^*/*ath1::MX*, during fermentations with cell recycling displayed slightly reduced cell viabilities, while maintaining ethanol titers similar to those of the reference, PE-2_H4 (S3 Fig).

We tested the double-mutant *cyr1^A1474T^*/*usv1Δ* by competitions against the tester-GFP via successive transfers in liquid YPS medium with 8% (v/v) ethanol and without ethanol (Fig 6C). At each passage the proportion of the competitors was recorded, and the cells were counted (S7 Table). Plotting the calculated cumulative S against the number of cell doublings at each passage allowed the estimation of S per doubling using linear regression obtained from the data points. Values were normalized for fitness calculated for the parental PE-2_H4 competing against tester-GFP. The double-mutant *cyr1^A1474T^*/*usv1Δ* exhibited a consistent fitness gain of 5.46% per doubling in the presence of 8% (v/v) ethanol, whereas it exhibited a fitness loss of 1.07% in the absence of ethanol (Fig 6C).

Finally, we subjected the engineered strain *cyr1^A1474T^*/*usv1Δ* and PE-2_H4 to five cycles simulating the sugarcane molasses fermentation process. A progressive increase (17.8%, 20.3%, 22.3%, 22.8%, and 22.5%) of total reducing sugars (TRS) from cycles 1 to 5 challenged the stress tolerance of the strains. Under this condition, the double-mutant *cyr1^A1474T^*/*usv1Δ* had accumulated approximately 1% more ethanol from cycles 3 to 5 than its parental (Fig 6D). More importantly, the average biomass gain of *cyr1^A1474T^*/*usv1Δ* was more than that of PE-2_H4 throughout the five fermentation cycles. Cell viability did not significantly differ between strains (Fig 6D).

The results from competition assays in 8% (v/v) ethanol indicated that mutations *cyr1^A1474T^*/*usv1Δ* improved strain ethanol tolerance in a rich medium. Furthermore, fermentations of sugarcane molasses with cell recycling by the double-mutant and P1c indicated fermentation performance with higher biomass formation (for *cyr1^A1474T^*/*usv1Δ*) and higher cell viabilities (for P1c) than that of the parental PE-2_H4. Although ethanol titers had not improved significantly, it should be noted that both strains had been challenged during fermentations via a combination of stress factors, such as toxic compounds in molasses, high ethanol levels, temperature, and acid treatment [10, 36]. Under these stresses, the *cyr1^A1474T^*/*usv1Δ* double-mutant and P1c exhibited higher capacities than their parental strains to sustain a viable biomass from one cycle to another. Such persistency represents the most important technological property for Brazilian bioethanol production as it allows the yeast to survive and maintain high ethanol production rates throughout the whole fermentation season of about eight months [10, 36, 52].

## DISCUSSION

Alcoholic tolerance is a key trait of yeasts used in large-scale production of the biofuel ethanol [1]. To understand the polygenic basis of alcohol tolerance for improving ethanologenic yeasts, several studies have been conducted using various approaches, such as transcriptomics [3, 53], screening of genome-wide knockout collections [53], global transcriptional machinery engineering [54], and QTL mapping [8, 12, 16]. Herein, we used ALE to investigate the genetic underpinnings of ethanol tolerance in the bioethanol strain PE-2_H4. Adaptive evolution has been used for raising the alcoholic tolerance of *S. cerevisiae* strains, mostly through protocols involving continuous propagation over hundreds of generations in the presence of increasing ethanol levels [15, 24–33]. Such studies highlighted the role of diploidization [15, 27], increase in cell size [31], and remodeling of membrane lipids [31] and cell wall [32] as important adaptations of yeasts to high ethanol concentrations.

Instead of propagation under constant ethanol exposure, our ALE approach, and that of two other studies [34, 35], favored cell survival to ethanol shocks as a paradigm for investigating ethanol tolerance in yeasts. The shock protocol that we applied was substantially different from these two previous ALE experiments, which used 10–12 rounds of quick 2-min exposures [34] or up to 30 cycles of 1–3 h shocks of up to 25% (v/v) ethanol [35]. These protocols also relied on a postshock propagation in rich culture medium to promote physiologic recovery and selection for growth fitness [34, 35]. Although, in those cases, the bimodal protocol supported ethanol tolerance combined with good fermentation performance [34, 35], in our ALE experiment, most reverse-engineered strains and evolved clones showed low growth fitness, either in the presence or absence of ethanol. Possibly, these side effects were observed only in our study because of the severe treatments that we used, which involved application of 2-hour ethanol exposures and usage of far more cycles of ethanol shocks (up to 82 rounds) with higher ethanol concentrations (up to 30% v/v) than those used before [34, 35].

To shed light on the genetic basis of cell survival during acute ethanol stress, our ALE study used WGS of evolved yeasts and performed a comprehensive evaluation of the fitness exhibited by reverse-engineered alleles. Regarding ALE protocols for ethanol tolerance in yeasts, genomic surveys were conducted in only two previous studies [15, 32]. One of them relied on three rounds of turbidostat cultivation with increasing amounts of ethanol. WGS of isolated clones revealed alleles associated with *SSD1* and *UTH1* as determinants of ethanol tolerance [32]. Another study was based on a turbidostat cultivation of six S288C populations for 200 generations with ethanol levels raised from 6% to 12% (v/v) [15]. The WGS uncovered hundreds of mutated genes associated with various functions, such as stress response, cell cycle control, DNA replication/repair, and respiration. Fitness measurements demonstrated that evolved alleles associated with *PRT1*, *MEX67*, and *VPS70* were adaptive to high levels of ethanol [15].

Similar to previous studies, our ALE experiment also detected diverse mutations (46 in total) related to various cellular functions, emphasizing the complex polygenic nature of ethanol tolerance trait in yeasts. However, our four evolved populations were particularly enriched in mutations affecting components of the cAMP/PKA signaling and trehalose degradation pathways. The Protein kinase A (PKA) complex, under glucose abundance, acts as an effector for cell proliferation, cell cycle progression, and ribosome biogenesis [39, 40]. Simultaneously, active PKA is an inhibitor of Msn2p and Msn4p transcription factors [37, 40, 41]. When glucose is depleted, causing low cAMP levels, the PKA-mediated inhibition is reverted and Msn2p and Msn4p become dephosphorylated and active; they shuttle into the nucleus to activate the environmental stress response, which includes factors involved in heat shock, cell wall remodeling, DNA repair, antioxidant defense, and trehalose biosynthesis [37, 40, 41]. Mutants that downregulate the Ras2p/cAMP pathway display a characteristically low PKA phenotype marked by the ectopic activation of Msn2/4p-mediated stress responses [37, 39–41]. In our ALE populations, we detected mutated alleles of *RAS2* and *CYR1*, as well as of *MDS3* and *PMD1*; knockouts of the latter two genes mimicked a low PKA phenotype [38]. We propose that downregulation of the PKA function is a central adaptation during ALE to mobilize yeast stress responses to withstand ethanol lethality. This was corroborated by the fact that evolved clones P1c, P2c, and P3c displayed higher trehalose accumulation and constitutive upregulation of *HSP12-GFP* expression, a standard biosensor responsive to Msn2/4p [37]. Accordingly, in our study, strains with the engineered *ras2::MX*, *mds3::MX*, and *cyr1^A1474T^* alleles exhibited higher fitness than the parental strain to tolerate the ethanol shock.

The disaccharide trehalose is a well-known stress protectant that possibly acts as a chemical chaperone to hold the folding of proteins and maintain the plasma membrane integrity during stress [55]. *ATH1* encodes an acid trehalase that is presumably involved in the extracellular degradation of trehalose [55, 56]. *Ath1* knockouts are tolerant to several stresses, such as heat [56], dehydration, freezing, and high ethanol concentration [57]. In three of our ALE populations, a*th1* defective alleles swept to dominance after a cAMP/PKA-related allele had emerged. Furthermore, we obtained the highest-fitness strain for tolerating ethanol shocks from the synergism between *cyr1^A1474T^* and *ath1::MX* mutations. Higher trehalose synthesis triggered by mutations that downregulate the cAMP/PKA pathways (e.g., *mds3*, *pmd1*, and *cyr1*) [38, 39] may potentiate the emergence of defective *ath1* alleles, blocking trehalose degradation and promoting its cellular accumulation and protective association with the plasma membrane [55–57].

Such peripheral protection may be an important mechanistic principle for ethanol tolerance, which is also supported by the snowflake phenotype of population P4. A defective allele affecting *BUD3* (acting on the axial bud site selection in haploid cells [48, 49]) probably compromises correct daughter cell separation following cytokinesis, resulting in multicellular aggregations. Possibly, within these clumps, inner cells may be shielded from ethanol damage, which is constrained to the cell cluster periphery [50]. Consistent with this idea is the fact that yeasts with aggregation phenotypes are more resistant to multiple stresses (including freeze/thaw, hydrogen peroxide, heat, and ethanol treatments) than individual cells [50, 51]. A further possible link between ethanol tolerance to cell wall shielding is provided by the *ROM2* alleles identified in P1 and P3. Rom2p is a guanine nucleotide exchange factor for Rho1p and Rho2p GTPases. It plays a role in the cell wall integrity signaling to remodel the cell wall in response to environmental stresses [58]. Moreover, Rom2p mediates stress resistance and cell growth by interacting with the Ras-cAMP pathway [59]. However, we could not demonstrate any fitness contribution to ethanol tolerance by the *rom2::MX* disruption or *rom2^G440R^* allele. A similar lack of validation is the case for alleles related to *PTR2*, which encodes a di–tripeptide transporter at the membrane [60]. Because we narrowed our reverse-engineering scope to a few genes or pathways that had parallel mutations in at least two populations, it is possible that *ROM2* and *PTR2* related alleles may be adaptive in association with mutations not tested in this study. Multiple genetic interactions and the diversity of 46 evolved alleles recovered via WGS in our ALE are consistent with the idea that ethanol tolerance may be achieved through various and complex mutational pathways [15].

Trade-off effects are common in ALE populations subjected to selection pressure for prolonged periods [17–19]. ALE populations tend to become specialists in the selective environment frequently displaying lower fitness than their ancestors when moved to an alternative niche, subjected to a different propagation mode, or nutrient condition [17–19]. Therefore, it is not surprising that several clones and reverse-engineered strains derived from yeast populations adapted to 68–82 ethanol shocks exhibit low growth fitness. We suggest that propagation fitness decay may partially reflect the overall downregulation of protein biogenesis and other growth-promoting components resulting from the constitutive activation of the environmental stress response [41, 61]. Slower growth rates related to mutants downregulating the PKA pathway and accumulating trehalose have been well documented [39–41]. The demonstrated inverse correlation between growth rates and resistance to severe stresses, a phenomenon that conforms to the principle of energy balance wherein resources used for growth under nonstress conditions are redirected to overcome the environmental stress [20–22], is relevant in our case, which involves yeasts tolerant to drastic ethanol treatments.

Considering the possible negative impact on fermentation performance, it is questionable whether strain selection protocols that apply harsh treatments are worth pursuing [17]. In our genetic analysis, we combined alleles *cyr1^A1474T^*/*usv1Δ* to generate a strain with higher fitness to acute ethanol treatments and better growth in the presence of ethanol than its parental strain. Usv1p is a C2H2 zinc finger transcription factor that activates gene expression under nonfermentable carbon sources and respiratory conditions [62, 63], and represses the transcription of genes involved in sulfur metabolism [64]. The fact that *USV1* takes part in transcriptional responses to hyperosmolarity and other stresses (including ethanol) [62, 65] seems to contradict that ethanol tolerance is conferred by the *usv1* knockout in the PE-2_H4 strain. Similarly, the reasons for positive epistasis of this mutation with *cyr1^A1474T^* are yet to be elucidated.

The alleles *usv1^Q73stop^* and *cyr1^A1474T^* are part of the P1c genetic makeup. Similar to the double-mutant *cyr1^A1474T^*/*usv1Δ*, P1c exhibited excellent performance under conditions simulating the Brazilian ethanol production system (i.e., using cell recycling and sulfuric acid shocks). These results indicate that harsh selection schemes may be useful for isolating strains suitable for bioprocesses wherein yeasts are subjected to multiple stresses; such as the fermentation of highly toxic biomass hydrolysates for cellulosic ethanol production [17]. Finally, even if shock-based screenings tend to present negative side effects, we suggest that it is possible to disentangle adaptive alleles from fitness-costing mutations through reverse engineering of adaptive alleles into a parental background (as shown here) [17, 19], or by applying sexual strategies to dissociate beneficial mutations from the deleterious ones [18, 66]. This may involve backcrossing shock-selected yeasts with their parental strains and subjecting the resulting recombinants to further selection for growth fitness under stress (de Bem and Gross, manuscript in preparation). These approaches may provide excellent complementary procedures for refining protocols for selection of stress-tolerant ethanologenic yeasts.

## MATERIALS AND METHODS

### Adaptive laboratory evolution

The *S. cerevisiae* haploid strains PE-2_H3 (*MATα*) and PE-2_H4 (*MATa*) are spore derivatives dissected from a tetrad of the Brazilian bioethanol isolate PE-2 [9, 36]. For populations P1, P2, and P3, a single progenitor was generated by integrating the natMX marker into the PE-2_H4 genome (S1 Table and S1 Text). Then, this transformant was mated with PE-2_H3 (*MATα*). After sporulation, a nourseothricin-resistant *MATα* spore was propagated to establish the P4 haploid population. All the strains used in this study are listed in S1 Table.

To initiate the ALE experiment, populations P1–P4 were propagated overnight in 20 mL of YPS medium (yeast extract-peptone plus 2% sucrose) in 50 mL Erlenmeyer flasks (60 rpm at 32°C). From this initial preinoculum, and after each recovery growth during ALE, 1 mL (containing approximately 10^8^ cells) was transferred into a 1.5 mL microcentrifuge tube and centrifuged at 8,000 rpm for 5 min. Then, the obtained pellet was washed once with 500 uL deionized water. Next, the cell pellet was resuspended in 1 mL PBS 1X buffer containing an initial ethanol concentration of 19% (v/v). During the ethanol treatment (i.e., the ethanol shock) cells were kept static at 32°C for 2 hours. Following treatment, cells were centrifuged at 8,000 rpm for 5 min, washed with 500 uL deionized water, resuspended, and transferred into 20 mL YPS for a recovery growth in a shaker incubator (32°C, 60 rpm). Nourseothricin (100 µg/mL) was supplemented to prevent contamination. The culture was maintained for 2–4 days, i.e., until the yeast cells achieved a stationary phase. Then, 1 mL of the culture was taken to begin a new shock/recovery cycle, and another 0.5 mL was cryopreserved in a 25% glycerol stock. Generally, during ALE, ethanol concentrations used for shocks were increased whenever the population showed fast postshock growth (e.g., reaching the stationary phase in 2 days). Conversely, the ethanol concentration was decreased whenever the postshock culture for a given population repeatedly showed prolonged recovery (approximately 4 days growth), indicating poor adaptation to the applied ethanol level. Final ethanol concentrations reached 30% (v/v) for populations P1 and P3, and 29% (v/v) for P2 and P4. Furthermore, the number of shock/recovery cycles varied among the populations (Fig 1B). At the end of the evolution, for each population, the single best growing colony on solid medium (YPS) supplemented with 8% (v/v) ethanol was selected for WGS and downstream analyses. These originated the evolved clones P1c, P2c, P3c, and P4c.

### WGS of evolved clones and variants identification

The evolved clones P1c–P4c were grown in liquid YPS until saturation. The sampled cellular mass from each clone was transferred into a lysis buffer (DNeasy® Plant Mini Kit, QIAGEN), and cells were disrupted by 1.5-min vortexing (Beadbeater, BioSpec) in the presence of zirconia beads. Genomic DNA was extracted from the lysate using the DNeasy® Plant Mini Kit (QIAGEN) following the manufacturer’s protocol. The isolated DNAs were fragmented using the NEBNext dsDNA Fragmentase (New England BioLabs), and paired-end libraries were constructed according to the Illumina TruSeq DNA PCR-Free Low Throughput Library Prep Kit (Illumina). The four paired-end libraries were quantified by qPCR using the KAPA Library Quantification Kit (Roche). Genome sequencing was conducted on the Illumina MiSeq platform at the Federal University of Rio Grande do Sul, Brazil, using the MiSeq Reagent Kit v3 supporting 600-cycles of 2 x 300 paired-end reads (Illumina). The resulting sequence reads were filtered following a cut-off of Phred quality scores ≥30 and read length ≥75 bases. The sequence reads obtained for P1c–P4c were submitted to NCBI (https://www.ncbi.nlm.nih.gov) under the BioProject number PRJNA1026594. Sequence reads derived from P1c, P2c, and P3c genomic libraries were mapped against the PE-2_H4 genome (GenBank accession number GCA_905220315.1) using the Burrows–Wheeler aligner algorithm implemented by the CLC Genomics Workbench 8.01 (QIAGEN). A cut-off of 0.8 for aligned read length and 0.8 for minimal required identity were set. The mapped reads were subjected to a variant detection performed on the CLC Genomics Workbench. A frequency of ≥50% was set as the cut-off; however, selected variants had frequencies tending to 100%, in accordance with a haploid background derived from a single colony. The progenitor of population P4 had a recombinant haploid genome derived from a PE-2_H4 vs. PE-2_H3 crossing. To identify mutations in the P4c sequence reads, we first mapped PE-2_H3 reads (GenBank accession number GCA_905220325.1) against the PE-2_H4 genome to obtain the coordinates for all polymorphisms (SNPs and small InDels) distinguishing the two genomes. We mapped the P4c sequence reads against the PE-2_H4 genome to call variants as described above. By comparing the coordinates obtained with those observed for the PE-2_H3 polymorphisms (Microsoft Excel), we identified P4c variants that were absent in PE-2_H3 and PE-2_H4 genomes. Key mapped mutations were confirmed by Sanger sequencing of PCR fragments derived from the final populations and their respective progenitors. PCR oligonucleotides were designed to flank the analyzed mutations (S4 Table). The presence of evolved alleles was also examined by Sanger sequencing of the PCR fragments derived from DNA extracted from cryopreserved intermediate populations (S1 Appendix).

### Yeast molecular genetics and strains construction

The strains constructed in this study were all derived from the Brazilian bioethanol yeast PE-2_H4 (*MATa*) [9] (S1 Table and S1 Text). An exception was the P4 progenitor (described above). Several strains were constructed with mutations mimicking the ALE evolved alleles. Constructs for insertional disruption of alleles via homologous recombination were assembled with PCR products of the targeted genes flanking the kanMX PCR fragment [42]. PCR reactions were performed using Phusion^®^ high-fidelity DNA polymerase according to the manufacturer’s protocol (New Englang BioLabs). Amplified flanking recombination regions were merged with the MX cassette through Circular Polymerase Exchange Cloning into the pUC19 plasmid [67]. Alternatively, a construct assembly was performed in two steps via in vivo cloning in *E. coli* [68]. Insertional kanMX cassettes were amplified using PCR (Phusion^®^ high-fidelity DNA polymerase), and about 500–1,000 ng PCR products were transformed into the PE-2_H4 via standard lithium acetate protocol [69]. In some cases, the kanMX cassette was directly amplified, with primers carrying tails with 40-nts homology at the 5′ region for integration into the targeted locus, and readily transformed into the yeast [42].

Targeted deletions and point mutations were introduced via CRISPR/Cas9 following the EasyGuide method developed by our group [43]. Donor sequences specifying genome edits were preassembled into the pUC19 vector, and then amplified using PCR (Phusion^®^ high-fidelity DNA polymerase), and cotransformed with gRNA-encoding PCR fragments for in vivo recombination [43]. Alternatively, donors were directly amplified by PCR with oligonucleotides carrying at least 40-nts homology arms for recombination, and readily cotransformed with gRNA-encoding PCR fragments [43]. Diagnostic PCRs (Taq DNA polymerase, Thermo Fisher Scientific) were routinely used to confirm genome edits and kanMX insertions. Single nucleotide edits were confirmed by Sanger sequencing. All primers used in this study for strain construction and authentication are listed in S4 Table. Molecular genetic procedures are detailed in S1 Text.

### Phenotypic analyses

Evolved clones P1c, P2c, and P3c were subjected to phenotypic analyses using the ALE progenitor as a reference. For cell viability, P1c–P3c and the progenitor were acclimatized by two shocks of ethanol 15% (v/v) and recovery growth. From the second outgrowth culture, 1 mL (10^8^ cells) was pelleted, washed, and resuspended in 1 mL PBS 1X buffer containing an ethanol concentration of 19% (v/v). Ethanol treatment lasted 2 h at 32°C. A 10^−5^ dilution was plated into solid YPS medium. After 3 days, colony forming units (CFU) were counted and compared to the number of CFUs obtained in a control treatment without ethanol (representing 100% survival). Assays were performed in triplicates.

Trehalose estimation following ethanol shocks was performed through the enzymatic trehalase assay [70]. Cells from P1c–P4c and the progenitor were acclimatized by two shocks of 15% (v/v) ethanol and outgrowths, according to our ALE protocol. About 10^8^ cells were taken from the last outgrowth, pelleted, washed, and resuspended in 1 mL PBS 1X buffer containing 19% (v/v) ethanol. The treatment lasted 2 hours before recovery growth in YPS. After 2 days, 10^8^ cells estimated on a Neubauer chamber were centrifuged and washed twice with cold water to remove any residual ethanol. The cells were resuspended in 0.25 M Na_2_CO_3_ solution, vortexed, and incubated at 95°C for about 3 hours. The pH was adjusted to 5.5, and the cells were vortexed and transferred to a fresh tube to estimate the amount of trehalose. Porcine kidney trehalase (Sigma-Aldrich) was used with a pH adjusted to 5.8. The cells were incubated at 37°C overnight. The glucose liberated was quantified using a glucose colorimetric detection kit, following the manufacturer’s protocol (Invitrogen) and normalized for the number of cells used in the sample. The amount of trehalose was calculated in µg of glucose-equivalents. Assays were conducted in triplicates.

The *HSP12-GFP* Msn2/4p-responsive biosensor was constructed by chromosomal integration (conducted using the CRISPR EasyGuide [43]) of the ORF expressing the ymUkG1 GFP [71] to produce an in frame fusion with the *HSP12* gene [37] (see S1 Text). The biosensor was constructed into the PE-2_H4 parental strain and P1c, P2c, P3c, and *cyr1^A1474T^* haploid backgrounds. The first assay was performed by propagating three replicates for each strain in YPS and recording fluorescence signals at the stationary and logarithmic growth phases over 4 days. Fluorescence measurements were obtained using the Attune NxT flow cytometer (Thermo Fisher Scientific) by recording 10,000 events. Fluorescence intensity values were taken from the median derived from flow cytometry histograms [37]. Fluorescence intensity values for the strains were normalized by the average numbers obtained for the PE-2_H4 reference at the same sampled point (Fig 3B). A second assay involved comparing fluorescence signals recorded from cells growing for 2 hours, 4 hours, 6 hours, 8 hours, and 24 hours after inoculation with or without 8% (v/v) ethanol. Normalization was made with the PE-2_H4 values at the same sampled points (Fig 3C). Alternatively, to demonstrate the effect of ethanol treatment on the *HSP12-GFP* expression for each strain, at each sample point values were expressed as a ratio of the fluorescence obtained for the ethanol exposed and nonexposed cells (Fig 3D).

### Competition experiments and fitness measurements

Haploid PE-2 yeasts tend to aggregate in groups of about two–six cells (Fig 5B). This mild aggregative phenotype is absent in the diploid background, when separate cells can be observed. To facilitate counting of individual cells by flow cytometry during competition assays, all strains used were diploidized by mating-type switching, induced via plasmidial expression of the HO endonuclease, followed by mating [72]. For an initial phenotypic evaluation, diploid P1c–P3, progenitor, and GFP-tagged PE- 2_H4 strain (tester-GFP, expressing the ymUkG1 GFP [71] integrated into the *HO* locus) were subjected to five passages in 20 mL YPS (with no addition of ethanol). Then, for each sample, about 5 × 10^7^ cells were mixed with an approximately equal amount (one : one) of the tester-GFP in a 1X PBS buffer. The initial proportion of GFP-marked and nontagged cells was then recorded (about 10,000 events in 10 µL) in the Attune NxT flow cytometer (Thermo Fisher Scientific) (S2 Table). Gating parameters were set with separate cultures of GFP-expressing cells and nonmarked yeasts. Mixed cells were distributed in three replicates, and each one was subjected to ethanol treatment (24% [v/v]) for 2 hours, and then to a recovery growth in 20 mL of YPS (32°C, 60 rpm) until the stationary phase was achieved. After outgrowth, the proportions of the competitors were estimated in the Attune NxT flow cytometer and plotted. In a parallel experiment, P1c–P3, progenitor, and tester-GFP were acclimatized by sequential 2-hour shocks of 15%, 15%, 20%, 20%, and 22% (v/v) ethanol, each one followed by an outgrowth. Then, ethanol-adapted cells were counted, mixed (one: one) with tester-GFP, and subjected to a 24% (v/v) ethanol treatment and recovery growth, as described above.

By recording the initial (i) percentage of GFP-marked (GFPi) and nonmarked evolved (EVOi) cells, and the final (f) proportion of competitors (GFPf and EVOf) after outgrowth, it is possible to calculate the selection coefficient (S), describing the fitness of the evolved cells, according to the following equation:

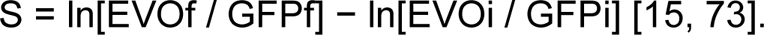

All selection coefficient values were normalized to the S obtained in a competition of the progenitor vs. tester-GFP (designated as S = 0). Therefore, S always expressed the fitness of evolved clones, or reverse-engineered strains, relative to the progenitor and discounted any fitness effects for GFP expression.

A slightly different ethanol shock assay was used for systematic analyses of evolved clones and reverse-engineered strains (Competition 1, Fig 4A and 4B). This included two cycles of adaptation with a 15% (v/v) ethanol shock and recovery, followed by a 20% (v/v) ethanol shock after mixing the evolved strain and the tester-GFP in an approximately one: one ratio (S5 Table). The proportion of competitors was recorded following outgrowth, and fitness was calculated using the equation above, using the S obtained for the ALE progenitor and parental PE-2_H4 strain used for reverse engineering for normalization. For competitions 2 and 3 (Fig 4A and 4B), the competitors were acclimatized via two passages in 20 mL YPS without ethanol and with 4% and 6% (v/v) ethanol, respectively. Competitors were then mixed with the tester-GFP in an approximately equal ratio. From this initial mixture, 20 µL for competition 2 and 100 µL for competition 3 were, respectively, inoculated in 20 mL for propagation at 32°C (60 rpm) up to the stationary phase. The proportion of GFP-labeled and nonlabeled cells were recorded by the end of the experiment. At the beginning and end of propagation, the cells were counted in the Attune NxT flow cytometer allowing the estimation of cell doublings during the propagation. A selection coefficient/cell doubling was estimated by dividing S for the total numbers of doublings (S/d) [15, 73]. All values for S/d were normalized to the data obtained for the parental strains (S5 Table).

Competitions with the *cyr1^A1474T^*/*usv1Δ* and PE-2_H4 against the tester-GFP were conducted through five passages in liquid YPS medium with 8% (v/v) ethanol or without ethanol. Acclimation and initiation of competitions were as described above. The proportion of competitors at the end of the first passage was stipulated as the starting point for measurements (S = 0). Cumulative S and cell doublings were estimated at each transfer (S7 Table). Values plotted for *cyr1^A1474T^*/*usv1Δ* were normalized for those obtained in the PE-2_H4 vs. tester-GFP competition. Selection coefficients per cell doublings (S/d) were calculated from the trendline fitted into the data points.

Microplate growth assays were conducted on the Tecan Sunrise© (Tecan). The preadaptation of strains with or without ethanol were as described above. About 2 x 10^6^ cells were inoculated into 200 µL medium per well. Assays were conducted in triplicates under static conditions at 28°C. The optical density (OD)600 was recorded every 15 min (S6 Table). Maximum specific growth rates (µmax) were calculated from OD values obtained during the exponential growth phase (ranging from OD 0.4 to 0.9). A plot of the natural logarithm of OD values (Microsoft Excel) against the collected time points (h^-1^) allowed fitting of a simple linear regression and obtaining the μ_max_ values from the slope [74]. The μmax was separately calculated for each replicate and expressed as an average value. At 8% (v/v) ethanol, strains *cyr1^A1434T^*, *ath1Δ*, *cyr1^A1434T^*/*ath1::MX*, and *usv1::MX* displayed a very poor growth profile, reaching a plateau of maximum optical density (OD_max_) below 0.9. In these cases, the µmax was obtained from ODs in the range between 0.1 and 0.5. The OD_max_ values were obtained at the stationary growth phase.

### Sugarcane molasses fermentation

Fermentations of sugarcane molasses were performed according to the scale down system proposed by Basso et al. [36] and described by Raghavendran et al. [10]. This closely simulates the Brazilian industrial fed-batch process with cell recycling and sulfuric acid treatment. Briefly, strains were propagated in molasses medium (10% w/v of the sugar hexose) at 32°C, centrifuged at 3,000 × g for 20 min and resuspended. For an initial inoculum, and at each new fermentation cycle, a cell suspension (70% w/v) was diluted in water up to 30% (12 mL) of the total fermentation volume (40 mL). The added substrate (diluted molasses) constituted the remaining 70% (28 mL), with TRS content adjusted to yield, at the end of each fermentation round, the intended ethanol titer, calculated according to ∼90% of the theoretical conversion rate of 0.511 g ethanol per g of reducing sugars (glucose/fructose) [10]. Fermentations were performed in triplicates at 34°C in a volume of 40 mL in 50 mL-centrifuge tubes. At the end of the fermentation, cell viability counts were taken as described by Basso et al. [36]. Cells were pelleted down before the wet biomass was weighed. Supernatant was saved for ethanol estimation as described by Basso et al. [36]. Preceding each cycle, collected biomass from the previous cycle was subjected to a sulfuric acid treatment, performed for 1 hour by adding 0.5 M H_2_SO_4_ (pH 2.5). Molasses feeding for the next cycle was performed as described above. Trehalose content was estimated from biomass collected at the beginning of the fermentations and at the end of the last cycles, as previously described by Basso et al. [36].

## SUPPORTING INFORMATION

**S1 Table. List of *Saccharomyces cerevisiae* strains.** Table listing all *S. cerevisiae* strains used in this study, including parental strains, evolved clones, and genetically-modified yeasts.

**S2 Table. Flow cytometry measurements in competitions with 24% v/v ethanol treatments.** The proportion of each competitor is shown before (initial) and after the 24% ethanol treatment and outgrowth (final). Selection coefficients were calculated (S) and normalized with the data obtained for the progenitor. Two datasets were generated: one for the Progenitor, P1c, P2c, P3c without acclimation, and other with the same strains subjected to prior acclimation with ethanol shocks (see text). The data was used to generate the graphics of Fig 1C and D.

**S3 Table. Results from WGS and variant calling of P1c, P2c, P3c, and P4c.** Mutations found in the evolved clones are shown. Coordinates are related to the PE- 2_H4 parental strain (GenBank accession number GCA_905220315.1).

**S4 Table. PCR oligonucleotides used in this study.** Oligonucleotides used for PCRs described in the S1 Text and S1 Appendix are listed.

**S5 Table. Flow cytometry measurements for competition assays 1–3.** Data from competitions 1 (green), 2 (red), and 3 (blue) display the proportions of tester-GFP and the indicated competitor at the initial and final points of the assay. Selection coefficients (S) were calculated and normalized with the S obtained for the respective parental strain. For competitions 2 and 3 the S was expressed per cell doubling.

**S6 Table. Microplate growth assay.** ODs were recorded every 15 min. on the Tecan Sunrise^TM^ microplate reader. Each Strain was assayed in 3–4 replicates in YPS with 8% (v/v) ethanol, or in YPS without ethanol.

**S7 Table. Fitness estimations for *cyr1^A1474T^*/*usv1Δ*.** The PE-2_H4 (parental) and *cyr1^A1474T^*/*usv1Δ* were competed against the PE-2_H4-GFP through five passages in YPS with 8% (v/v) ethanol or without ethanol. Flow cytometry records of the competitors’ proportions and cell counting were taken at the initial and final points of each transfer, allowing the calculation of selection coefficients (S) during each passage and a cumulative S during serial transfers 2–5. By plotting the cumulative S per number of calculated generations at the end of each passage, selective coefficients per cell doublings (S/d) were calculated from the trendline fitted into the data points. Thereby an S per doubling was calculated for PE-2_H4 as -0.0079 for the competition with the PE-2_H4-GFP at 8% (v/v) ethanol and as 0.0049 for the competition in YPS without ethanol. Based on these numbers, a “S correction factor for PE-2_H4” was calculated accounting for cumulative cell doublings at each passage and used to normalize the cumulative S obtained for *cyr1^A1474T^*/*usv1Δ* in a competition with PE-2_H4_GFP. This normalization allows to express the *cyr1^A1474T^*/*usv1Δ* fitness (S) in comparison to PE-2_H4 and to discount any fitness effect of GFP expression in the PE-2_H4-GFP strain. The normalized cumulative S for *cyr1^A1474T^*/*usv1Δ* was plotted in Fig 6C.

**S1 Text. Strain construction procedures.** Molecular cloning and molecular genetics approaches used during construction of *S. cerevisiae strains* used in this study are explained.

**S1 Appendix. Sanger sequencing chromatograms for the wild-type and evolved alleles.** Sanger sequencing allowed identification of wild-type and evolved alleles for each population (P1–P4), according to the ethanol shock/recovery cycles. Alleles are indicated by red arrows. Two chromatogram peaks are observed in cycles where the wild-type and evolved alleles coexist in a population. Primers pairs used for PCR amplifications for each sequencing reaction are indicated.

**S1 Fig. Median fluorescence intensities in strains expressing the *HSP12-GFP* biosensor.** Median values of florescence intensities obtained for strains PE-2_H4, P1c, P2c, P3c and *cyr1^A1474T^* expressing *HSP12-GFP* are shown. These data were used to generate the heatmaps in Fig 3 of the main text. (**A**) As in Fig 3B, the time course of GFP fluorescence signal during four consecutive passages (four days) in YPS (without ethanol) for strains is shown. Median florescence values were obtained at the stationary (*sta*) and logarithmic (*log*) growth phases. (**B**) The same data as in (A) expressed as fluorescence fold changes relative to the PE-2_H4 signal at the same time point. (**C**, **D**) Time course of GFP fluorescence along 24 hrs in cells propagating without ethanol (C) and in 8% (v/v) ethanol (D). The data from (C and D) was the basis for the heatmap depicted in Fig 3C and D of the main text. Statistical analyses refer to the mutant strain being compared to the PE-2_H4 at the same time point. (*) p<0.05, one way ANOVA followed by Bonferroni post-test for multiple comparisons.

**S2 Fig. The flocculation phenotype of *bud3* disruption depends on the haploid state.** Only *bud3::MX* haploid cells (left) exhibited flocculation. Diploid *bud3::MX* cells (right) were no longer aggregated. The diploid state was confirmed by PCR of the *MAT* locus showing the two mating-types.

**S3 Fig. Ethanol production and cell viability of engineered strains during fermentations.** Reverse-engineered strains *ath1Δ*, *cyr1^A1474T^*, and *cyr1^A1474T^*/*ath1::MX* were compared with the parental PE-2_H4 through 14 cycles of sugarcane molasse fermentations. (**A**) At each new cycle, total reducing sugars concentrations were progressively raised to increase the percentage of ethanol production (v/v). Overall, ethanol production performance of genetically-modified strains was not better than the parental PE-2_H4. (**B**) Higher ethanol levels decreased the cell viability of tested yeasts. Generally, genetically-modified strains were more sensitive to the ethanol levels than the parental PE-2_H4.

## Notes

### Competing Interest Statement

The authors have declared no competing interest.

## REFERENCES

1. Jacobus AP, Gross J, Evans JH, Ceccato-Antonini SR, Gombert AK. Saccharomyces cerevisiae strains used industrially for bioethanol production. Essays Biochem. 2021;65(2):147–61. Epub 2021/06/23. doi: 10.1042/EBC20200160.

2. Piskur J, Rozpedowska E, Polakova S, Merico A, Compagno C. How did Saccharomyces evolve to become a good brewer? Trends Genet. 2006;22(4):183–6. Epub 2006/02/28. doi: 10.1016/j.tig.2006.02.002.

3. Ma M, Liu ZL. Mechanisms of ethanol tolerance in Saccharomyces cerevisiae. Appl Microbiol Biotechnol. 2010;87(3):829–45. Epub 2010/05/14. doi: 10.1007/s00253-010-2594-3.

4. Ding J, Huang X, Zhang L, Zhao N, Yang D, Zhang K. Tolerance and stress response to ethanol in the yeast Saccharomyces cerevisiae. Appl Microbiol Biotechnol. 2009;85(2):253–63. Epub 2009/09/17. doi: 10.1007/s00253-009-2223-1.

5. Yang KM, Lee NR, Woo JM, Choi W, Zimmermann M, Blank LM, et al. Ethanol reduces mitochondrial membrane integrity and thereby impacts carbon metabolism of Saccharomyces cerevisiae. FEMS Yeast Res. 2012;12(6):675–84. Epub 2012/06/16. doi: 10.1111/j.1567-1364.2012.00818.x.

6. Voordeckers K, Colding C, Grasso L, Pardo B, Hoes L, Kominek J, et al. Ethanol exposure increases mutation rate through error-prone polymerases. Nat Commun. 2020;11(1):3664. Epub 2020/07/23. doi: 10.1038/s41467-020-17447-3.

7. Snoek T, Picca Nicolino M, Van den Bremt S, Mertens S, Saels V, Verplaetse A, et al. Large-scale robot-assisted genome shuffling yields industrial Saccharomyces cerevisiae yeasts with increased ethanol tolerance. Biotechnol Biofuels. 2015;8:32. Epub 2015/03/12. doi: 10.1186/s13068-015-0216-0.

8. Swinnen S, Schaerlaekens K, Pais T, Claesen J, Hubmann G, Yang Y, et al. Identification of novel causative genes determining the complex trait of high ethanol tolerance in yeast using pooled-segregant whole-genome sequence analysis. Genome Res. 2012;22(5):975–84. Epub 2012/03/09. doi: 10.1101/gr.131698.111.

9. Jacobus AP, Stephens TG, Youssef P, Gonzalez-Pech R, Ciccotosto-Camp MM, Dougan KE, et al. Comparative Genomics Supports That Brazilian Bioethanol Saccharomyces cerevisiae Comprise a Unified Group of Domesticated Strains Related to Cachaca Spirit Yeasts. Front Microbiol. 2021;12:644089. Epub 2021/05/04. doi: 10.3389/fmicb.2021.644089.

10. Raghavendran V, Basso TP, da Silva JB, Basso LC, Gombert AK. A simple scaled down system to mimic the industrial production of first generation fuel ethanol in Brazil. Antonie Van Leeuwenhoek. 2017;110(7):971–83. Epub 2017/05/05. doi: 10.1007/s10482-017-0868-9.

11. Della-Bianca BE, Basso TO, Stambuk BU, Basso LC, Gombert AK. What do we know about the yeast strains from the Brazilian fuel ethanol industry? Appl Microbiol Biotechnol. 2013;97(3):979–91. Epub 2012/12/29. doi: 10.1007/s00253-012-4631-x.

12. Pais TM, Foulquie-Moreno MR, Hubmann G, Duitama J, Swinnen S, Goovaerts A, et al. Comparative polygenic analysis of maximal ethanol accumulation capacity and tolerance to high ethanol levels of cell proliferation in yeast. PLoS Genet. 2013;9(6):e1003548. Epub 2013/06/12. doi: 10.1371/journal.pgen.1003548.

13. Snoek T, Verstrepen KJ, Voordeckers K. How do yeast cells become tolerant to high ethanol concentrations? Curr Genet. 2016;62(3):475–80. Epub 2016/01/14. doi: 10.1007/s00294-015-0561-3.

14. Casey GP, Ingledew WM. Ethanol tolerance in yeasts. Crit Rev Microbiol. 1986;13(3):219–80. Epub 1986/01/01. doi: 10.3109/10408418609108739.

15. Voordeckers K, Kominek J, Das A, Espinosa-Cantu A, De Maeyer D, Arslan A, et al. Adaptation to High Ethanol Reveals Complex Evolutionary Pathways. PLoS Genet. 2015;11(11):e1005635. Epub 2015/11/07. doi: 10.1371/journal.pgen.1005635.

16. Duitama J, Sanchez-Rodriguez A, Goovaerts A, Pulido-Tamayo S, Hubmann G, Foulquie-Moreno MR, et al. Improved linkage analysis of Quantitative Trait Loci using bulk segregants unveils a novel determinant of high ethanol tolerance in yeast. BMC Genomics. 2014;15:207. Epub 2014/03/20. doi: 10.1186/1471-2164-15-207.

17. Menegon YA, Gross J, Jacobus AP. How adaptive laboratory evolution can boost yeast tolerance to lignocellulosic hydrolyses. Curr Genet. 2022;68(3-4):319–42. Epub 2022/04/02. doi: 10.1007/s00294-022-01237-z.

18. Barrick JE, Lenski RE. Genome dynamics during experimental evolution. Nat Rev Genet. 2013;14(12):827–39. Epub 2013/10/30. doi: 10.1038/nrg3564.

19. Dragosits M, Mattanovich D. Adaptive laboratory evolution -- principles and applications for biotechnology. Microb Cell Fact. 2013;12:64. Epub 2013/07/03. doi: 10.1186/1475-2859-12-64.

20. Levy SF, Ziv N, Siegal ML. Bet hedging in yeast by heterogeneous, age-correlated expression of a stress protectant. PLoS Biol. 2012;10(5):e1001325. Epub 2012/05/17. doi: 10.1371/journal.pbio.1001325.

21. Ho YH, Gasch AP. Exploiting the yeast stress-activated signaling network to inform on stress biology and disease signaling. Curr Genet. 2015;61(4):503–11. Epub 2015/05/11. doi: 10.1007/s00294-015-0491-0.

22. Zakrzewska A, van Eikenhorst G, Burggraaff JE, Vis DJ, Hoefsloot H, Delneri D, et al. Genome-wide analysis of yeast stress survival and tolerance acquisition to analyze the central trade-off between growth rate and cellular robustness. Mol Biol Cell. 2011;22(22):4435–46. Epub 2011/10/04. doi: 10.1091/mbc.E10-08-0721.

23. Varize CS, Bücker A, Lopes LD, Christofoleti-Furlan RM, Raposo MS, Basso LC, et al. Increasing Ethanol Tolerance and Ethanol Production in an Industrial Fuel Ethanol Saccharomyces cerevisiae Strain. Fermentation. 2022;8(10: 470). doi: 10.3390/fermentation8100470.

24. Brown SW, Oliver SG. Isolation of ethanol-tolerant mutants of yeast by continuous selection. Appl Microbiol Biotechnol. 1982;16:119–22. doi: 10.1007/BF00500738.

25. Zazulyaa A, Semkiva M, Dmytruka K, Sibirnya A. Adaptive Evolution for the Improvement of Ethanol Production During Alcoholic Fermentation with the Industrial Strains of Yeast Saccharomyces cerevisiae. Cytology and Genetics. 2020;54(5):398–407. doi: 10.3103/S0095452720050059.

26. Fiedurek J, Skowronek M, Gromada A. Selection and adaptation of Saccharomyces cerevisae to increased ethanol tolerance and production. Pol J Microbiol. 2011;60(1):51–8. Epub 2011/06/03.

27. Turanli-Yildiz B, Benbadis L, Alkim C, Sezgin T, Aksit A, Gokce A, et al. In vivo evolutionary engineering for ethanol-tolerance of Saccharomyces cerevisiae haploid cells triggers diploidization. J Biosci Bioeng. 2017;124(3):309–18. Epub 2017/05/30. doi: 10.1016/j.jbiosc.2017.04.012.

28. Zhang M, Zhu R, Zhang M, Wang S. Creation of an ethanol-tolerant Saccharomyces cerevisiae strain by 266 nm laser radiation and repetitive cultivation. J Biosci Bioeng. 2014;118(5):508–13. Epub 2014/06/25. doi: 10.1016/j.jbiosc.2014.04.016.

29. M. N, R. G, E. B, M. M, M. Y, Morales P. Improved fermentation kinetics by wine yeast strains evolved under ethanol stress. Food Sci Technol. 2014;58:166–72. doi: 10.1016/j.lwt.2014.03.004.

30. Stanley D, Fraser S, Chambers PJ, Rogers P, Stanley GA. Generation and characterisation of stable ethanol-tolerant mutants of Saccharomyces cerevisiae. J Ind Microbiol Biotechnol. 2010;37(2):139–49. Epub 2009/11/11. doi: 10.1007/s10295-009-0655-3.

31. Dinh TN, Nagahisa K, Hirasawa T, Furusawa C, Shimizu H. Adaptation of Saccharomyces cerevisiae cells to high ethanol concentration and changes in fatty acid composition of membrane and cell size. PLoS One. 2008;3(7):e2623. Epub 2008/07/10. doi: 10.1371/journal.pone.0002623.

32. Avrahami-Moyal L, Engelberg D, Wenger JW, Sherlock G, Braun S. Turbidostat culture of Saccharomyces cerevisiae W303-1A under selective pressure elicited by ethanol selects for mutations in SSD1 and UTH1. FEMS Yeast Res. 2012;12(5):521–33. Epub 2012/03/27. doi: 10.1111/j.1567-1364.2012.00803.x.

33. Jimenez J, Benitez T. Selection of Ethanol-Tolerant Yeast Hybrids in pH-Regulated Continuous Culture. Appl Environ Microbiol. 1988;54(4):917–22. Epub 1988/04/01. doi: 10.1128/aem.54.4.917-922.1988.

34. Yang J, Tavazoie S. Regulatory and evolutionary adaptation of yeast to acute lethal ethanol stress. PLoS One. 2020;15(11):e0239528. Epub 2020/11/11. doi: 10.1371/journal.pone.0239528.

35. Mavrommati M, Papanikolaou S, Aggelis G. Improving ethanol tolerance of Saccharomyces cerevisiae through adaptive laboratory evolution using high ethanol concentrations as a selective pressure. Process Biochemistry. 2023;124:280–9. doi: 10.1016/j.procbio.2022.11.027.

36. Basso LC, de Amorim HV, de Oliveira AJ, Lopes ML. Yeast selection for fuel ethanol production in Brazil. FEMS Yeast Res. 2008;8(7):1155–63. Epub 2008/08/30. doi: 10.1111/j.1567-1364.2008.00428.x.

37. Sadeh A, Movshovich N, Volokh M, Gheber L, Aharoni A. Fine-tuning of the Msn2/4- mediated yeast stress responses as revealed by systematic deletion of Msn2/4 partners. Mol Biol Cell. 2011;22(17):3127–38. Epub 2011/07/16. doi: 10.1091/mbc.E10-12-1007.

38. McDonald CM, Wagner M, Dunham MJ, Shin ME, Ahmed NT, Winter E. The Ras/cAMP pathway and the CDK-like kinase Ime2 regulate the MAPK Smk1 and spore morphogenesis in Saccharomyces cerevisiae. Genetics. 2009;181(2):511–23. Epub 2008/12/18. doi: 10.1534/genetics.108.098434.

39. Van Dijck P, Ma P, Versele M, Gorwa MF, Colombo S, Lemaire K, et al. A baker’s yeast mutant (fil1) with a specific, partially inactivating mutation in adenylate cyclase maintains a high stress resistance during active fermentation and growth. J Mol Microbiol Biotechnol. 2000;2(4):521–30.

40. Plank M. Interaction of TOR and PKA Signaling in S. cerevisiae. Biomolecules. 2022;12(2). Epub 2022/02/26. doi: 10.3390/biom12020210.

41. Smith A, Ward MP, Garrett S. Yeast PKA represses Msn2p/Msn4p-dependent gene expression to regulate growth, stress response and glycogen accumulation. EMBO J. 1998;17(13):3556–64. Epub 1998/07/03. doi: 10.1093/emboj/17.13.3556.

42. Guldener U, Heck S, Fielder T, Beinhauer J, Hegemann JH. A new efficient gene disruption cassette for repeated use in budding yeast. Nucleic Acids Res. 1996;24(13):2519–24. Epub 1996/07/01. doi: 10.1093/nar/24.13.2519.

43. Jacobus AP, Barreto JA, de Bem LS, Menegon YA, Fier I, Bueno JGR, et al. EasyGuide Plasmids Support in Vivo Assembly of gRNAs for CRISPR/Cas9 Applications in Saccharomyces cerevisiae. ACS Synth Biol. 2022;11(11):3886–91. Epub 2022/10/19. doi: 10.1021/acssynbio.2c00348.

44. Ratcliff WC, Denison RF, Borrello M, Travisano M. Experimental evolution of multicellularity. Proc Natl Acad Sci U S A. 2012;109(5):1595–600. Epub 2012/02/07. doi: 10.1073/pnas.1115323109.

45. Opalek M, Wloch-Salamon D. Aspects of Multicellularity in Saccharomyces cerevisiae Yeast: A Review of Evolutionary and Physiological Mechanisms. Genes (Basel). 2020;11(6). Epub 2020/07/01. doi: 10.3390/genes11060690.

46. Oud B, Guadalupe-Medina V, Nijkamp JF, de Ridder D, Pronk JT, van Maris AJ, et al. Genome duplication and mutations in ACE2 cause multicellular, fast-sedimenting phenotypes in evolved Saccharomyces cerevisiae. Proc Natl Acad Sci U S A. 2013;110(45):E4223–31. Epub 2013/10/23. doi: 10.1073/pnas.1305949110.

47. Rodrigues-Prause A, Sampaio NMV, Gurol TM, Aguirre GM, Sedam HNC, Chapman MJ, et al. A Case Study of Genomic Instability in an Industrial Strain of Saccharomyces cerevisiae. G3 (Bethesda). 2018;8(11):3703–13. Epub 2018/09/27. doi: 10.1534/g3.118.200446.

48. Sanders SL, Field CM. Cell division. Bud-site selection is only skin deep. Curr Biol. 1995;5(11):1213–5. Epub 1995/11/01. doi: 10.1016/s0960-9822(95)00239-9.

49. Kang PJ, Lee ME, Park HO. Bud3 activates Cdc42 to establish a proper growth site in budding yeast. J Cell Biol. 2014;206(1):19–28. Epub 2014/07/09. doi: 10.1083/jcb.201402040.

50. Kuzdzal-Fick JJ, Chen L, Balazsi G. Disadvantages and benefits of evolved unicellularity versus multicellularity in budding yeast. Ecol Evol. 2019;9(15):8509–23. Epub 2019/08/15. doi: 10.1002/ece3.5322.

51. Alugoju P, Mahilkar A, Saini S. Evolution of multicellularity and unicellularity in yeast S. cerevisiae to study reversibility of evolutionary trajectories. bioRxiv. 2020;2020.08.15.252361. doi: 10.1101/2020.08.15.252361.

52. Rego-Costa A, Huang IT, Desai MM, Gombert AK. Yeast population dynamics in Brazilian bioethanol production. G3 (Bethesda). 2023;13(7). Epub 2023/06/02. doi: 10.1093/g3journal/jkad104.

53. Stanley D, Bandara A, Fraser S, Chambers PJ, Stanley GA. The ethanol stress response and ethanol tolerance of Saccharomyces cerevisiae. J Appl Microbiol. 2010;109(1):13–24. Epub 2010/01/15. doi: 10.1111/j.1365-2672.2009.04657.x.

54. Alper H, Moxley J, Nevoigt E, Fink GR, Stephanopoulos G. Engineering yeast transcription machinery for improved ethanol tolerance and production. Science. 2006;314(5805):1565-8. Epub 2006/12/13. doi: 10.1126/science.1131969.

55. Eleutherio E, Panek A, De Mesquita JF, Trevisol E, Magalhaes R. Revisiting yeast trehalose metabolism. Curr Genet. 2015;61(3):263–74. Epub 2014/09/12. doi: 10.1007/s00294-014-0450-1.

56. Magalhaes RSS, Popova B, Braus GH, Outeiro TF, Eleutherio ECA. The trehalose protective mechanism during thermal stress in Saccharomyces cerevisiae: the roles of Ath1 and Agt1. FEMS Yeast Res. 2018;18(6). Epub 2018/07/15. doi: 10.1093/femsyr/foy066.

57. Kim J, Alizadeh P, Harding T, Hefner-Gravink A, Klionsky DJ. Disruption of the yeast ATH1 gene confers better survival after dehydration, freezing, and ethanol shock: potential commercial applications. Appl Environ Microbiol. 1996;62(5):1563–9. Epub 1996/05/01. doi: 10.1128/aem.62.5.1563-1569.1996.

58. Levin DE. Regulation of cell wall biogenesis in Saccharomyces cerevisiae: the cell wall integrity signaling pathway. Genetics. 2011;189(4):1145–75. Epub 2011/12/17. doi: 10.1534/genetics.111.128264.

59. Park JI, Collinson EJ, Grant CM, Dawes IW. Rom2p, the Rho1 GTP/GDP exchange factor of Saccharomyces cerevisiae, can mediate stress responses via the Ras-cAMP pathway. J Biol Chem. 2005;280(4):2529–35. Epub 2004/11/17. doi: 10.1074/jbc.M407900200.

60. Hauser M, Narita V, Donhardt AM, Naider F, Becker JM. Multiplicity and regulation of genes encoding peptide transporters in Saccharomyces cerevisiae. Mol Membr Biol. 2001;18(1):105–12. Epub 2001/06/09.

61. Gasch AP, Werner-Washburne M. The genomics of yeast responses to environmental stress and starvation. Funct Integr Genomics. 2002;2(4-5):181–92. Epub 2002/08/23. doi: 10.1007/s10142-002-0058-2.

62. Hlynialuk C, Schierholtz R, Vernooy A, van der Merwe G. Nsf1/Ypl230w participates in transcriptional activation during non-fermentative growth and in response to salt stress in Saccharomyces cerevisiae. Microbiology (Reading). 2008;154(Pt 8):2482–91. Epub 2008/08/01. doi: 10.1099/mic.0.2008/019976-0.

63. Sameith K, Amini S, Groot Koerkamp MJ, van Leenen D, Brok M, Brabers N, et al. A high-resolution gene expression atlas of epistasis between gene-specific transcription factors exposes potential mechanisms for genetic interactions. BMC Biol. 2015;13:112. Epub 2015/12/25. doi: 10.1186/s12915-015-0222-5.

64. Bessonov K, Walkey CJ, Shelp BJ, van Vuuren HJ, Chiu D, van der Merwe G. Functional analyses of NSF1 in wine yeast using interconnected correlation clustering and molecular analyses. PLoS One. 2013;8(10):e77192. Epub 2013/10/17. doi: 10.1371/journal.pone.0077192.

65. Costa ACT, Russo M, Fernandes AAR, Broach JR, Fernandes PMB. Transcriptional Response of Multi-Stress-Tolerant Saccharomyces cerevisiae to Sequential Stresses. Fermentation. 2023;9(195). doi: 10.3390/fermentation9020195.

66. McDonald MJ, Rice DP, Desai MM. Sex speeds adaptation by altering the dynamics of molecular evolution. Nature. 2016;531(7593):233-6. Epub 2016/02/26. doi: 10.1038/nature17143.

67. Quan J, Tian J. Circular polymerase extension cloning of complex gene libraries and pathways. PLoS One. 2009;4(7):e6441. Epub 2009/08/04. doi: 10.1371/journal.pone.0006441.

68. Jacobus AP, Gross J. Optimal cloning of PCR fragments by homologous recombination in Escherichia coli. PLoS One. 2015;10(3):e0119221. Epub 2015/03/17. doi: 10.1371/journal.pone.0119221.

69. Gietz RD, Woods RA. Transformation of yeast by lithium acetate/single-stranded carrier DNA/polyethylene glycol method. Methods Enzymol. 2002;350:87–96. Epub 2002/06/21. doi: 10.1016/s0076-6879(02)50957-5.

70. Parrou JL, Francois J. A simplified procedure for a rapid and reliable assay of both glycogen and trehalose in whole yeast cells. Anal Biochem. 1997;248(1):186–8. Epub 1997/05/15. doi: 10.1006/abio.1997.2138.

71. Kaishima M, Ishii J, Matsuno T, Fukuda N, Kondo A. Expression of varied GFPs in Saccharomyces cerevisiae: codon optimization yields stronger than expected expression and fluorescence intensity. Sci Rep. 2016;6:35932. Epub 2016/10/27. doi: 10.1038/srep35932.

72. Herskowitz I, Jensen RE. Putting the HO gene to work: practical uses for mating-type switching. Methods Enzymol. 1991;194:132–46. Epub 1991/01/01. doi: 10.1016/0076-6879(91)94011-z.

73. Galeota-Sprung B, Guindon B, Sniegowski P. The fitness cost of mismatch repair mutators in Saccharomyces cerevisiae: partitioning the mutational load. Heredity (Edinb). 2020;124(1):50–61. Epub 2019/09/14. doi: 10.1038/s41437-019-0267-2.

74. Rodrigues CIS, Della-Bianca BE, Gombert AK. μMAX of Saccharomyces Cerevisiae: So Often Used, So Seldom Put into Perspective. Research Square. 2021. doi: 10.21203/rs.3.rs-182823/v1.

