## Supporting Figures for "Optimal trade-off between boosted tolerance and growth fitness during adaptive evolution of yeast to ethanol shocks"

### Supporting Figures S1–S3

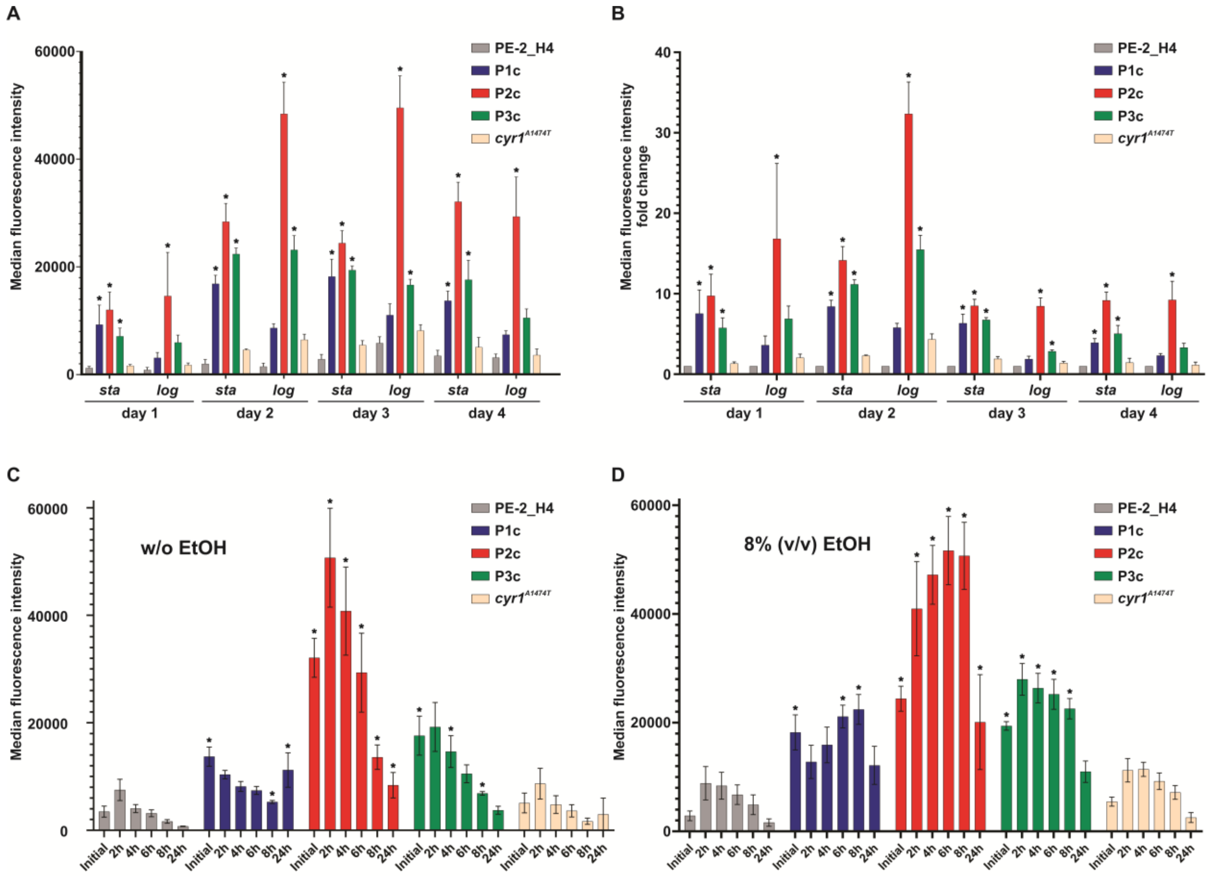


**S1 Fig. Median fluorescence intensities in strains expressing the *HSP12-GFP* biosensor.** Median values of florescence intensities obtained for strains PE-2_H4, P1c, P2c, P3c and *cyr1*$^{A1474T}$ expressing *HSP12-GFP* are shown. These data were used to generate the heatmaps in Fig 3 of the main text. (**A**) As in Fig 3B, the time course of GFP fluorescence signal during four consecutive passages (four days) in YPS (without ethanol) for strains is shown. Median florescence values were obtained at the stationary (*sta*) and logarithmic (*log*) growth phases. (**B**) The same data as in (A) expressed as fluorescence fold changes relative to the PE-2_H4 signal at the same time point. (**C**, **D**) Time course of GFP fluorescence along 24 hrs in cells propagating without ethanol (C) and in 8% (v/v) ethanol (D). The data from (C and D) was the basis for the heatmap depicted in Fig 3C and D of the main text. Statistical analyses refer to the mutant strain being compared to the PE-2_H4 at the same time point. (*) $p<0.05$, one way ANOVA followed by Bonferroni post-test for multiple comparisons.

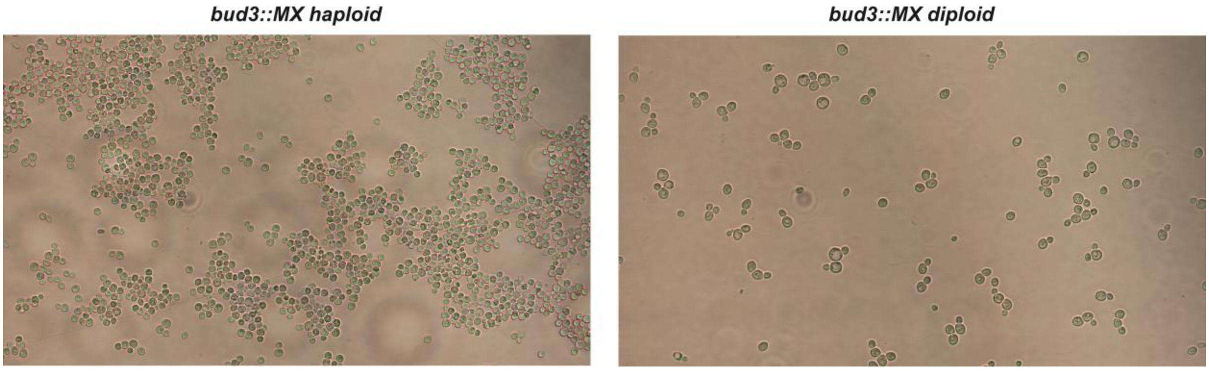


**S2 Fig. The flocculation phenotype of *bud3* disruption depends on the haploid state.** Only *bud3::MX* haploid cells (left) exhibited flocculation. Diploid *bud3::MX* cells (right) were no longer aggregated. The diploid state was confirmed by PCR of the *MAT* locus showing the two mating-types.

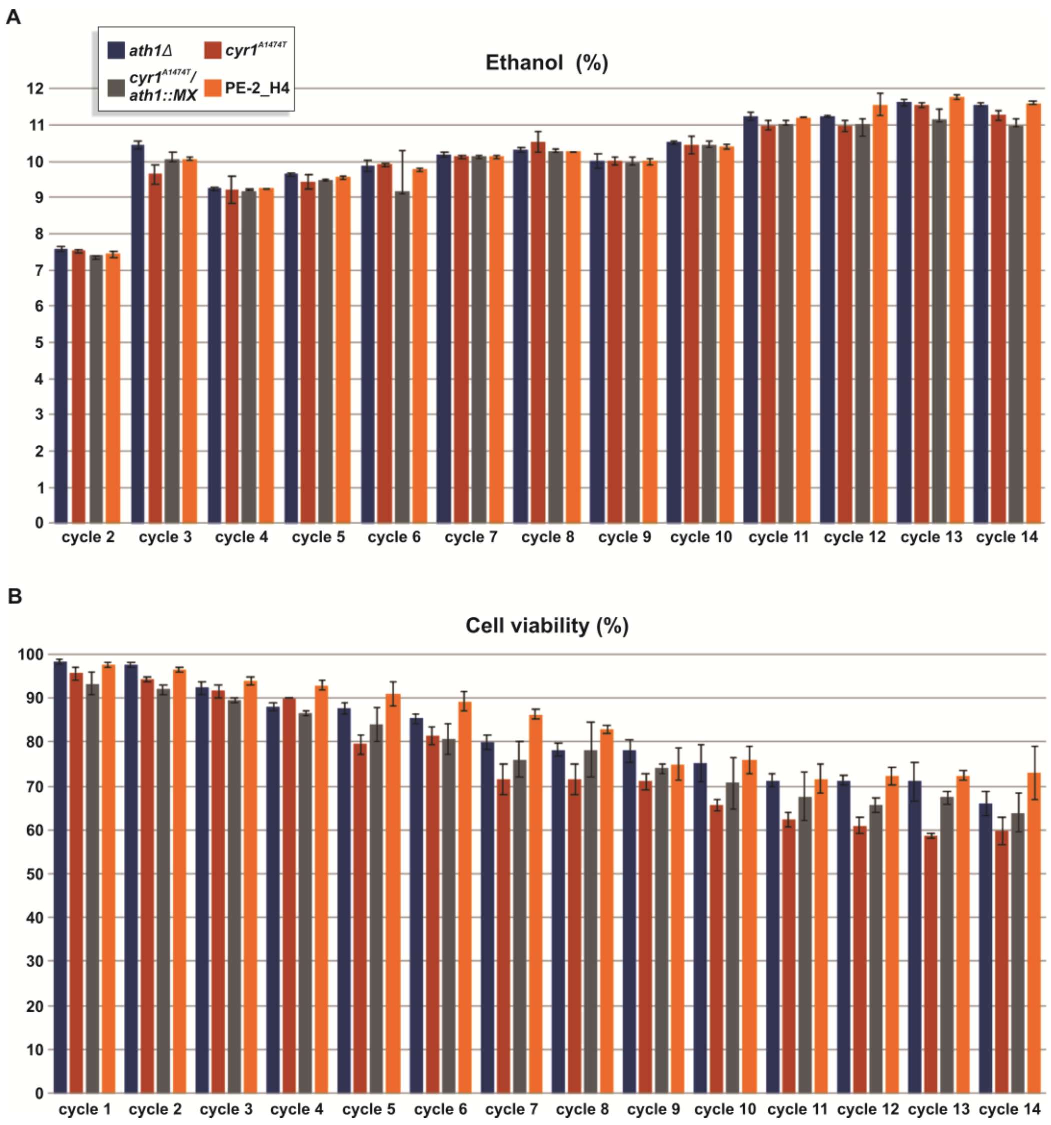


**S3 Fig. Ethanol production and cell viability of engineered strains during fermentations.** Reverse-engineered strains *ath1Δ*, *cyr1*$^{A1474T}$, and *cyr1*$^{A1474T}$/*ath1::MX* were compared with the parental PE-2_H4 through 14 cycles of sugarcane molasse fermentations. (**A**) At each new cycle, total reducing sugars concentrations were progressively raised to increase the percentage of ethanol production (v/v). Overall, ethanol production performance of genetically-modified strains was not better than the parental PE-2_H4. (**B**) Higher ethanol levels decreased the cell viability of tested yeasts. Generally, genetically-modified strains were more sensitive to the ethanol levels than the parental PE-2_H4.
