## Supplementary material for "Optimal trade-off between boosted tolerance and growth fitness during adaptive evolution of yeast to ethanol shocks": S1 Text

**S1 Text. Strain construction procedures**

### STRAIN CONSTRUCTION PROCEDURES

Molecular cloning procedures for plasmid construction in *E. coli* followed the methods described for in vivo cloning in *E. coli* [1] and the Circular Polymerase Extension Cloning (CPEC) [2]. Both are homology-based PCR cloning methods relying on oligonucleotides designed to have ≥20 nucleotides (nts) of overlap between joined PCR fragments and the vector backbone. Oligonucleotides used as PCR primers were synthetized at Exxtend Biotechnology (Campinas, Brazil) and are listed in S4 Table. CRISPR procedures followed our previously published CRISPR EasyGuide approach, relying on in vivo homologous recombination in *S. cerevisiae* of PCR-amplified plasmid parts that recombine to form functional gRNAs [3]. Genetic transformation in *S. cerevisiae* followed the lithium acetate protocol [4]. The antibiotics concentrations used to select transformed colonies and to propagate strains were: 200 µg/mL geneticin, 250 µg/mL zeocin, 100 µg/mL nourseothricin, and 300 µg/mL hygromycin. PCRs for cloning and genetic modifications were performed using the Phusion® high-fidelity DNA polymerase (New England BioLabs), and diagnostic PCRs using the Taq DNA polymerase (Thermo Fisher Scientific). The use of these enzymes was according to the manufacturer’s instructions.

#### Construction of parental strains for competition assays and ALE

Mating-type switching and diploidization of the PE-2_H4 *MATa* strain [5] (and other PE-2_H4 *MATa* haploid derived strains used in this work) was carried out via plasmid-based expression of the HO endonuclease [6]. We cloned a functional *HO* gene via in vivo cloning in *E. coli* [1]. For that, a *HO* PCR product, amplified with primers HOXhoI and HOHindIII, was cloned in fusion with the *GAL1* promoter contained in the pSH65 plasmid backbone opened with the XhoI/HindIII restriction enzymes [7]. The resulting pSHO was transformed into the PE_H4 haploid strain and selected with zeocin. For diploidization, individual transformed colonies were inoculated in YP-galactose 2% for overnight growth and induction of the *HO* expression. Next day, stationary grown cells were plated into YPD 2% solid medium.

Mating-type of the resulting colonies was probed by PCR with the primers *MAT*, *MATa*, and *MATalpha* as described in reference [8]. Diploid cells were selected. In the case where diploids could not be isolated, and cells of the opposite *MATα*-type were identified, the latter were grown and mixed with the *MATa* cells in liquid medium for mating and diploidization.

For integration of the kanMX marker into the PE-2_H4 parental, the *HO* locus was amplified from the genome of this strain using PCR with the primers HoLf and HoRr. The resulting 896 bp DNA fragment was cloned via in vivo recombination in *E. coli* [1] into the pUC19 vector [9], which was amplified by PCR with primers PUClonERfor and PUClonERrev (2,288 bp). The resulting pUCHO plasmid was then linearized via PCR (2,288 bp) with the HoLr and HoRf oligonucleotides, treated with DpnI (New England Biolabs), and column-purified (QIAquick PCR purification kit, QIAGEN). For cloning, the linearized vector was cotransformed into *E. coli* with a 1,439 bp KanMX PCR fragment amplified with primers KanpB3for and KanpB3rev from the pUG6 plasmid [10]. In vivo recombination generated the pHOkan plasmid. PCR conditions and in vivo cloning procedures were as described in reference [1]. The cassette for chromosomal integration into the *HO* locus was amplified from the pHOkan using PCR with primers HoLf and HoRr and transformed into the PE-2_H4 strain. Geneticin-resistant colonies were propagated for DNA extraction. PCRs with this genomic DNA confirmed the cassette integration by using primers HoEf/ProMX_rev (649 bp product for the left integration site) and HoEr/TermMX_for (604 bp product for the right integration site). The resulting H4::*MX* strain was subjected to diploidization as described above.

To generate the PE-2_H4 tester-GFP strain, a guide RNA (gRNA) was designed to target the *HO* locus via the CRISPR EasyGuide system [3]. For in vivo yeast assembly of two gRNAs specifying the target site, the pEasyG3-zeo and pEasyG3-mic backbones were amplified by PCR with primers gA_HO and gB_HO. In parallel, two split donor sequences were generated by PCR to assembly one GFP-expressing construct via recombination and chromosome integration into the *HO* locus. One PCR product of 409 bp, corresponding to the promoter *pTEF1*, was amplified from pUG6 [10] with primers TefGFP_f/TefGFP_r. The PCR *GFP-termTDH1* fragment of 1,050 bp was generated with primers GFP_f/TDH1_r from the pRS426_GFP plasmid (expressing the ymUkG1 GFP [11]). This plasmid was a gift from Dr. Leandro V. dos

Santos (University of Campinas, Brazil). Both split donor fragments were co-transformed with the gRNA parts into the PE-2_H4 strain expressing the pEasyCas9 plasmid. Integration and correct construct assembly were confirmed by PCR with primers HoEf/ProMX_rev (211 bp on the left flank) and TDH1term_f/ HoRr (251 bp on the right flank), and by detection of the GFP fluorescence on the Attune NxT flow cytometer (Thermo Fisher Scientific). The resulting strain was diploidized as described above.

The *HSP12-GFP* biosensors were assembled into the PE-2_H4, P1c, P2c, P3c, and *cyr1*$^{A1474T}$ haploid backgrounds. For that, a gRNA was designed to target the *HSP12* CDS at its 3' region via CRISPR EasyGuide. For in vivo yeast assembly of two gRNAs specifying the target site, the pEasyG3-hph and pEasyG3-mic backbones were amplified by PCR with primers gA_HSP12 and gB_HSP12. A donor DNA of 767 bp was PCR-amplified from the pRS426_GFP plasmid with primers H12_GFPf2/ H12_GFPr. This donor and the PCR-amplified plasmid parts for gRNA assembly were cotransformed into strains carrying the pEasyCas9 plasmid. Homology arms specified by the 5' regions of the donors ensured homologous recombination of *GFP* in frame with the *HSP12* gene. Correct integration was confirmed by PCR with primers H12_f/GFPcheck_r (239 bp) and GFP_f2/ H12ter_r (249 bp).

The progenitor for ALE populations P1, P2, and P3 was tagged with the nourseothricin resistance marker by integration of the plasmid pUC19-natMX into the rRNA-encoding cluster on Chr. XII of PE-2_H4. We previously cloned the pUCnat plasmid by inserting the natMX cassette into the pUC19 vector [1]. The pUCnat was linearized by PCR with primers pUCrDNArev/ MXrRNA_f. The 3,986 bp product was treated with DpnI and purified (QIAquick PCR purification kit, QIAGEN). To amplify the rRNA DNA target region, two overlapping PCR fragments were generated from the PE-2_H4 DNA with primers rDNA1for/rDNANotrev (1,008 bp) and rDNANotfor/rDNA1rev (1,128 bp). A NotI restriction site was placed into the PCR oligonucleotides within a 50-nts overlapping region connecting the two fragments. Both PCR products were merged with the pUCnat fragment via CPEC [2]. The resulting plasmid pUC19-natMX combined the natMX cassette with 2,087 bp of the rRNA-encoding region spanning the non-translated region NTS1-2 and the 5' part of the 25S ribosomal RNA. The pUC19-natMX was linearized by cleavage with NotI (New England Biolabs) to facilitate its chromosomal integration via homologous recombination. After the transformation and selection procedures, the presence of the pUC19-natMX in the genomic DNA was confirmed by PCR with primers Termfor/pUC19-2rev (149 bp) and by the stable segregation of the nourseothricin resistance in transformants submitted to serial culturing. The P4 *MATα* progenitor resulted from a cross between the P1-P3 progenitor and the PE-2_H3 strain [5].

**Gene disruptions via kanMX insertions**

For disruptions of *RAS2*, *MDS3*, *ATH1*, and *USV1* ORFs, insertion cassettes containing homologous integration regions flanking the KanMX cassette [10] were pre-assembled into the *E. coli* pUC19 plasmid. For that, the KanMX cassette was PCR-amplified with the KanpB3for/KanpB3rev primers (1,439 bp), and the pUC19 backbone was linearized by PCR with primers PUClonERfor/PUClonERrev (2,288 bp). Left and right flanking regions for each target were respectively amplified by PCR from the PE-2_H4 genome with the following primer combinations: *RAS2*: Ras2Lf/Ras2Lr (558 bp) and Ras2Rf/Ras2Rr (588 bp); *MDS3*: Mds3Lf/Mds3Lr (506 bp) and Mds3Rf/Mds3Rr (533 bp); *ATH1*: Ath1Lf/Ath1Lr (479 bp) and Ath1Rf/Ath1Rr (481 bp); *USV1*: Usv1Lf/Usv1Lr (548 bp) and Usv1Rf/Usv1Rr (660 bp). Flanking PCR fragments were fused with the KanMX cassette and pUC19 linearized backbone in a CPEC reaction. Plasmid-assembled integration cassettes were amplified by PCR with left and right primers as follows: *RAS2*: Ras2Lf/Ras2Rr (2,537 bp); *MDS3*: Mds3Lf/Mds3Rr (2,428 bp); *ATH1*: Ath1Lf/Ath1Rr (2,349 bp); *USV1*: Usv1Lf/Usv1Rr (2,599 bp). Amplified products were transformed into the PE-2_H4 strain via the lithium acetate protocol [4] and selected with geneticin (200 µg/mL). Correct integrations were confirmed by diagnostic PCRs of transformants with the following primers combinations: *RAS2*: RAS2diaF/ProMX_rev (199 bp) and TermMX_for/RAS2diaR (262 bp); *MDS3*: MDS3diaF/ProMX_rev (266 bp) and TermMX_for/MDS3diaR (151 bp); *ATH1*: ATH1diaF/ProMX_rev (267 bp) and TermMX_for/ATH1diaR (137 bp); *USV1*: USV1diaF/ProMX_rev (199 bp) and TermMX_for/USV1diaR (195 bp). The strains were diploidized for their use in competition assays.

For *PTR2*, *ROM2* and *BUD3* disruptions, the KanMX cassette was directly PCR-amplified from the pUG6 with primers containing about 40 bp homology arms to recombine with the target regions [10]. Primers combinations were as follows: *PTR2*: PTR2MXfor/PTR2MXrev (1,519 bp); *ROM2*: ROM2MX-f/ROM2MX-r (1,519 bp); and *BUD3*: BUD3MX_f/BUD3MX_r (1,519 bp). After transformation and selection, diagnostic colony PCRs confirmed the insertions with the following primer combinations: *PTR2*: PTR2for/ProMX_rev (208 bp) and TermMX_for/PTR2P4rev2 (206 bp); *ROM2*: ROM2P1f2/ProMX_rev (238 bp) and TermMX_for/ROM2P1r2 (261 bp); *BUD3*: BUD3ExtL/ProMX_rev (232 bp) and TermMX_for/BUD3ExtR (281 bp). The strains were diploidized for their use in competition assays.

**Genome editing of SNPs and deletions via CRISPR EasyGuide**

The gRNAs for CRISPR-mediated genetic modifications were assembled according to the CRISPR EasyGuide method [3]. The 20-nts gRNA spacer sequences were predicted using CRISPOR [12]. The selected spacer sequences were specified at the 5’ regions of forward and reverse primers gA and gB, respectively, used for amplification of pEasyG3 modules and for in vivo assembly of functional gRNAs [3]. The gA and gB oligonucleotides for each targeted locus are listed in S4 Table.

The *PTR2* (T1084G) point mutation (strain *PTR2*$^{W362G}$) was introduced by a donor composed of two 70-nts oligos (DonPTR2-f and DonPTR2-r) having 22 nts of complementarity at their 3’ regions. The two oligos were hybridized and submitted to 35 cycles of extension PCR to form a 118 bp double-stranded donor DNA [13]. The overlapping region between the oligos specified the mutation T1084G in the *PTR2* CDS (Trp362Gly), and further neutral modifications in the PAM and gRNA recognition regions (the encoded amino acids were not changed) [14]. A KpnI cutting site, present in the WT sequence, was also modified to facilitate the identification of DNA-edited transformants. The 22 bp overlapping region was flanked at both sides by 48 bp providing homology arms for recombination into the *PTR2* locus. The double-stranded donor was cotransformed with the PCR-amplified gRNA fragments (pEasyG3) into the PE-2_H4 strain expressing the Cas9. Transformants were first selected by generating from their genomes a 256 bp PCR product with primers PTR2_f2/PTR2_r1. The DNA fragment was subjected to a restriction analysis with KpnI (New England Biolabs), and the failure to cut the 256 bp PCR product indicated possible genome-edited sites. Following this trial, the genome edition was confirmed by Sanger sequencing of a 590 bp PCR product amplified with primers PTR2for/PTR2rev from the genome of pre-selected transformants.

For generating the *rom2*$^{G440R}$ strain, a donor DNA was precloned into *E. coli*. The pUC19 was amplified by PCR with primers Vf/Vr to generate a linear vector of 1,725 bp. Two overlapping PCR fragments were generated from the *ROM2* locus with primer combinations DonLf_ROM2f/DonLr_ROM2f (329 bp) and DonRf_ROM2f/ DonRr_ROM2f (378 bp). The overlapping region of 30 bp contained the G1318A (Gly440Arg) mutation, and specified modifications of the PAM and gRNA recognition regions, without changing the encoded amino acids. A SalI cutting site was also introduced to facilitate genetic identification. The donor was assembled into the pUC19 vector by CPEC [2]. For the CRISPR experiment, the donor was amplified using PCR with primers ROM2fDon_f/ROM2fDon_r (653 bp) and cotransformed with the pEasyG3 PCR fragments into the PE-2_H4 carrying the pEasyCas9. To screen the transformants, the cleavage of a 266 bp PCR product (ROM2P3for/ROM2P3_r1) with SalI (New England Biolabs) into two fragments (82 bp and 184 bp) indicated a possible successful genetic modification. For confirmation, a 909 bp PCR product was generated from the genomic DNA with primers ROM2f_Seq1/ROM2f_Seq2. Sanger sequencing of this PCR product with primers ROM2f_Seq1, ROM2f_Seq2, ROM2P3for, and ROM2P3rev validated the genetic modifications. The $rom2^{G440R}$ modification was also introduced into the $cyr1^{A1474T}$ background.

The preassembly of a donor sequence into *E. coli* was also used to specify the mutations for strain $cyr1^{A1474T}$ construction. Three PCR fragments were generated from *CYR1* for the donor assembly. The PCR1 of *CYR1*, with primers CYRDonor2Lf/CYRDonor2Lr, generated a 233 bp fragment. The PCR2 amplified a 173 bp product with primers CYRDonor2Mf/CYRDonor2Mr, and the PCR3 generated a 302 bp fragment with primers CYRDonor2Rf/CYRDonor2Rr. PCR products 1, 2, and 3 had short overlapping sequences allowing the connection between adjacent fragments and to the pUC19 backbone. In a CPEC procedure [2], the PCR products 1, 2, and 3 were fused to the pUC19 vector backbone linearized via PCR with primers PUClonERfor/pUClonerRev (2,288 bp). A 609 bp donor was amplified from the resulting pDON2Cyr1 via PCR with primers Don2Cyr1f/CYR1r2. Two gRNA targets were selected for *CYR1*. Following the CRISPR EasyGuide protocol [3], spacers for gRNAs 1 and 2 were specified by amplifying via PCR the pEasyG2-mic, with primers gA1_CYR1/gB2_CYR2, and the pEasyG2-zeo, with oligos gA2_CYR1/gB1_CYR1 (S4 Table). The two pEasyG2-amplified modules recombined in vivo in yeast to assemble two functional gRNAs [3]. In the preassembled donor sequence, the overlapping sequence of 21 nts between PCR products 1 and 2 specified neutral mutations obliterating the recognition of the gRNA1, whereas the 27 nts shared between PCRs2 and 3 modified the site for gRNA2 recognition and introduced the encoded amino acid substitution Ala1474Thr. The donor contained flanking regions of 186 bp left and 261 bp right, allowing its homologous recombination into the *CYR1* locus. Transformation of donor and gRNA DNAs into the PE-2_H4 expressing the Cas9 generated the intended genetic modifications and introduced primer-binding sites for PCR screening. A PCR amplification with primers 2NestCYR1f/2NestCYR1r generates a 173 bp product only in the *cyr1*$^{A1474T}$ strain, whereas a PCR with primers CYR1test_f/CYR1test_r generates a product of 163 bp only in the WT PE-2_H4 strain. For confirmation of mutations, a PCR product of 753 bp, spanning the modified genomic region, was generated with primers DON2Cyr1Ef/DON2Cyr1Er. The same primers were used for sequencing the PCR product via the Sanger method.

The *ras2Δ* was constructed into the *ptr2*$^{W362G}$ background. For that, a donor DNA of 179 bp was generated by PCR amplification of the genomic locus with primers DonRAS2_f/Ras2Err. The bipartite primer DonRAS2_f was designed to specify a 1,012 bp deletion of the *RAS2* gene; i.e., its 5' region contained 40 nts homology to the left flank of the deletion, whereas the 20 nts at the 3' region annealed 1,012 bp apart, delimiting the deletion's right flank. After transformation via CRISPR EasyGuide, confirmation of the deletion was possible by a colony-PCR with primers Ras2Elf/Ras2Err, generating a 199 bp product. A PCR product of 275 bp with primers RAS2diaF/RAS2diaR corresponded to the *RAS2* WT locus. The *ath1Δ* strain was also constructed by producing a 173 bp donor DNA via PCR with the bipartite forward primer DonATH1_f and the reverse primer ATH1Er. After transformation, confirmation of the 2,038 bp deletion was possible by a colony-PCR with primers ATHP2for/ATH1Er, generating a 206 bp product. A PCR product of 218 bp with primers ATH1diaF/ATH1diaR corresponded to the *ATH1* WT locus. For construction of *mds3Δ* mutation into the *cyr1*$^{A1474T}$ and *rom2*$^{G440R}$ backgrounds, a DNA donor of 182 bp was generated via PCR with a bipartite forward primer DonMDS3_f and the reverse oligonucleotide Mds3Er. The CRISPR/Cas9 mutagenesis with this donor generated a 1,421 bp deletion of *MDS3*. This was confirmed by a PCR with primers MDS3P3for/Mds3Er, generating a 266 bp fragment. Amplification of the intact WT *MDS3* with primers MDS3diaF/MDS3diaR produces a 231 bp fragment. Finally, the *usv1Δ* strain was generated into *cyr1*$^{A1474T}$ background by the same strategy. A donor was amplified from the genome with the primers DonUSV1_f/ USV1Er (175 bp). Following CRISPR/Cas9 mutagenesis, the 1,106 bp *USV1* deletion was confirmed by

PCR with primers USV1Ef/USV1Er, generating a product of 203 bp. A PCR with primers USV1diaF/USV1diaR of the WT *USV1* locus produced a fragment of 208 bp. All strain produced via CRISPR EasyGuide were diploidized to facilitate their use in competition assays and flow cytometry analysis.

### References


1. Jacobus AP, Gross J. Optimal cloning of PCR fragments by homologous recombination in Escherichia coli. PLoS One. 2015;10(3):e0119221. Epub 2015/03/17. doi: 10.1371/journal.pone.0119221.
2. Quan J, Tian J. Circular polymerase extension cloning of complex gene libraries and pathways. PLoS One. 2009;4(7):e6441. Epub 2009/08/04. doi: 10.1371/journal.pone.0006441.
3. Jacobus AP, Barreto JA, de Bem LS, Menegon YA, Fier I, Bueno JGR, et al. EasyGuide Plasmids Support in Vivo Assembly of gRNAs for CRISPR/Cas9 Applications in Saccharomyces cerevisiae. ACS Synth Biol. 2022;11(11):3886-91. Epub 2022/10/19. doi: 10.1021/acssynbio.2c00348.
4. Gietz RD, Woods RA. Transformation of yeast by lithium acetate/single-stranded carrier DNA/polyethylene glycol method. Methods Enzymol. 2002;350:87-96. Epub 2002/06/21. doi: 10.1016/s0076-6879(02)50957-5.
5. Jacobus AP, Stephens TG, Youssef P, Gonzalez-Pech R, Ciccotosto-Camp MM, Dougan KE, et al. Comparative Genomics Supports That Brazilian Bioethanol Saccharomyces cerevisiae Comprise a Unified Group of Domesticated Strains Related to Cachaca Spirit Yeasts. Front Microbiol. 2021;12:644089. Epub 2021/05/04. doi: 10.3389/fmicb.2021.644089.
6. Herskowitz I, Jensen RE. Putting the HO gene to work: practical uses for mating-type switching. Methods Enzymol. 1991;194:132-46. Epub 1991/01/01. doi: 10.1016/0076-6879(91)94011-z.
7. Gueldener U, Heinisch J, Koehler GJ, Voss D, Hegemann JH. A second set of loxP marker cassettes for Cre-mediated multiple gene knockouts in budding yeast. Nucleic Acids Res. 2002;30(6):e23. Epub 2002/03/09. doi: 10.1093/nar/30.6.e23.
8. Huxley C, Green ED, Dunham I. Rapid assessment of S. cerevisiae mating type by PCR. Trends Genet. 1990;6(8):236. Epub 1990/08/01. doi: 10.1016/0168-9525(90)90190-h.
9. Norrander J, Kempe T, Messing J. Construction of improved M13 vectors using oligodeoxynucleotide-directed mutagenesis. Gene. 1983;26(1):101-6. Epub 1983/12/01. doi: 10.1016/0378-1119(83)90040-9.
10. Guldener U, Heck S, Fielder T, Beinhauer J, Hegemann JH. A new efficient gene disruption cassette for repeated use in budding yeast. Nucleic Acids Res. 1996;24(13):2519-24. Epub 1996/07/01. doi: 10.1093/nar/24.13.2519.
11. Kaishima M, Ishii J, Matsuno T, Fukuda N, Kondo A. Expression of varied GFPs in Saccharomyces cerevisiae: codon optimization yields stronger than expected expression and fluorescence intensity. Sci Rep. 2016;6:35932. Epub 2016/10/27. doi: 10.1038/srep35932.
12. Concordet JP, Haeussler M. CRISPOR: intuitive guide selection for CRISPR/Cas9 genome editing experiments and screens. Nucleic Acids Res. 2018;46(W1):W242-W5. Epub 2018/05/16. doi: 10.1093/nar/gky354.
13. Zhang Y, Wang J, Wang Z, Zhang Y, Shi S, Nielsen J, et al. A gRNA-tRNA array for CRISPR-Cas9 based rapid multiplexed genome editing in Saccharomyces cerevisiae. Nat Commun. 2019;10(1):1053. Epub 2019/03/07. doi: 10.1038/s41467-019-09005-3.

14. Horwitz AA, Walter JM, Schubert MG, Kung SH, Hawkins K, Platt DM, et al. Efficient Multiplexed Integration of Synergistic Alleles and Metabolic Pathways in Yeasts via CRISPR-Cas. Cell Syst. 2015;1(1):88-96. Epub 2016/05/03. doi: 10.1016/j.cels.2015.02.001.
