## Supplementary material for "Optimal trade-off between boosted tolerance and growth fitness during adaptive evolution of yeast to ethanol shocks": S1 Appendix

**S1 Appendix. Sanger sequencing chromatograms for the wild-type and evolved alleles.**

#### POPULATION 1 (P1)

##### *CYR1* (G4420A) (Ala1474Thr)

**PCR: Cyr1P1for/Cyr1P1rev; sequencing: Cyr1P1for**

**Progenitor**

**4420 (G)**

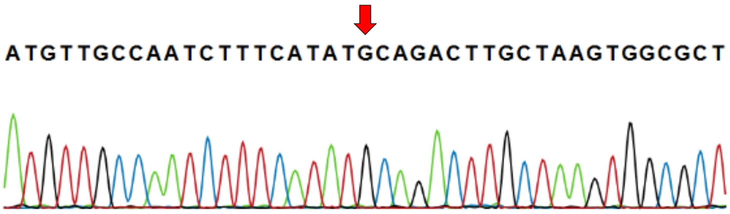


**Cycle 20**

**4420 (A)**

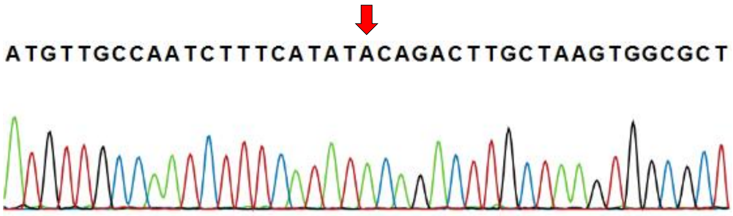


##### *MDS3* (1589InsG) (Val530fs)

**PCR: MDS3P1for/MDS3P1rev; sequencing: MDS3P1for**

**Progenitor**

**1589 (T)**

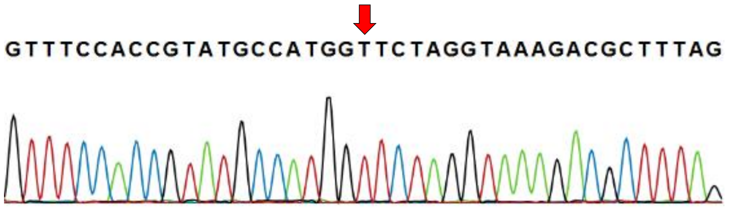


**Cycle 20**

**1589 (G)**

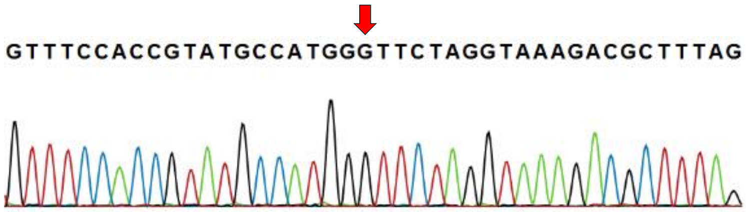


##### *ROM2* (C866T) (Ser289Leu)

**PCR: ROM2P1for/ROM2P1rev; sequencing: ROM2P1for**

**Progenitor**

**866 (C)**

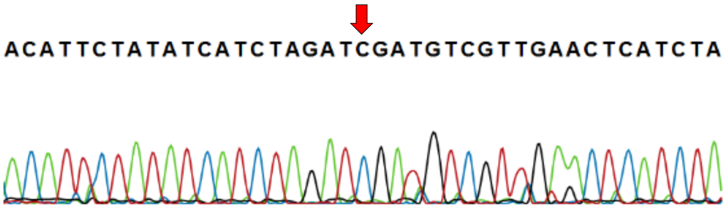


**Cycle 20**

**866 (T)**

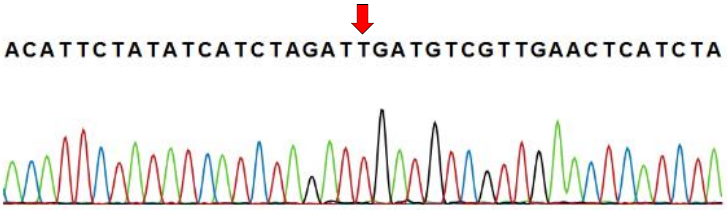

#### *ATH1* (1863DelT) (Phe621fs)

**PCR: ATH1P1for/ATH1P1rev; sequencing: ATH1P1for**

**Progenitor**

**1863 (T)**

ATCAAAAACGCAGCCCAATTTTTTACTGCTTATGTTAAGTA

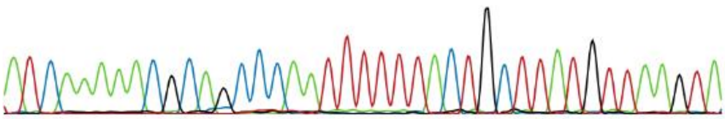

**Cycle 40**

**1863 (A/T)**

ATCAAAAACGCAGCCCAATTTTTACTGGTTATGKTAARKAC

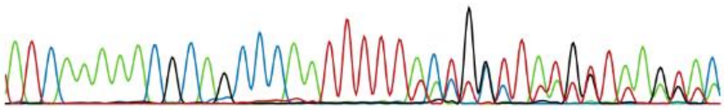

**Cycle 60**

**1863 (A/T)**

ATCAAAAACGCAGCCCAATTTTTACTGSTTATGTTAAGTAC

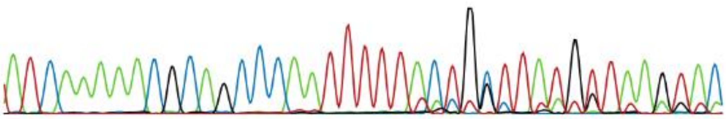

**Cycle 68 (end)**

**1863 (A)**

ATCAAAAACGCAGCCCAATTTTTACTGCTTATGTTAAGTAC

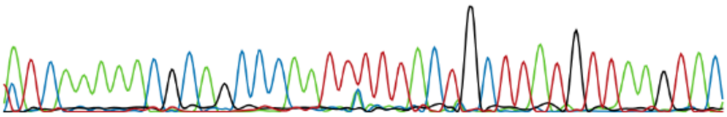

#### *USV1* (C217T) (Gln73Stop)

**PCR: USV1for/USV1rev; sequencing: USV1for**

**Cycle 20**

**217 (C)**

ACACTGGTGAAAAGCCGTTCCAGTGTCCTGCATGTTTGAAA

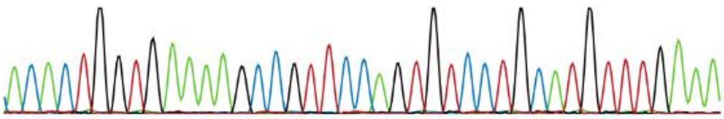

**Cycle 40**

**217 (T)**

ACACTGGTGAAAAGCCGTTCTAGTGTCCTGCATGTTTGAAA

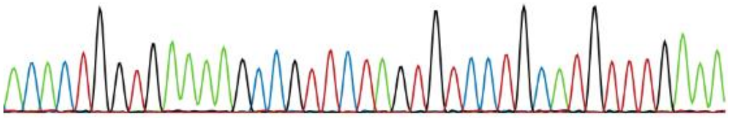

#### *DIG1* (C753A) (Tyr251Stop)

PCR: DIG1P1for/DIG1P1rev; sequencing DIG1P1for

Cycle 60

753 (C)

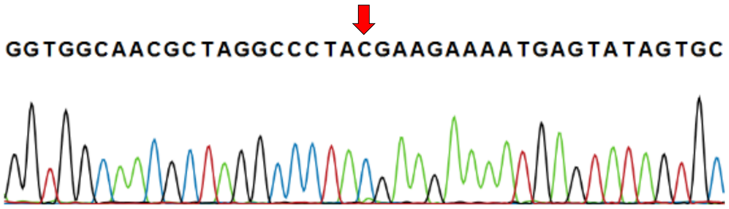


Cycle 68 (end)

753 (A)

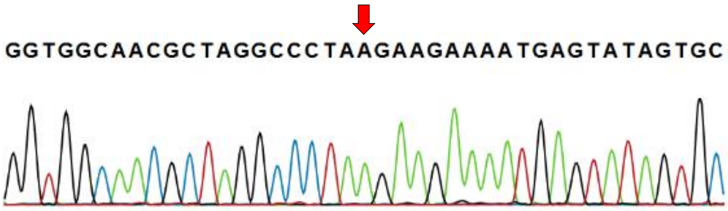


#### POPULATION 2 (P2)

#### *CYR1* (G2763T) (Leu921Phe)

PCR: CYR1P2for/CYR1P2rev; sequencing: CYR1P2for

Progenitor

2763 (G)

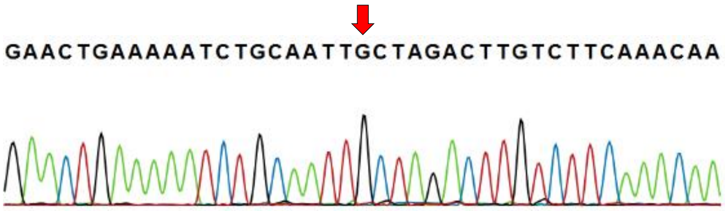


Cycle 20

2763 (T/G)

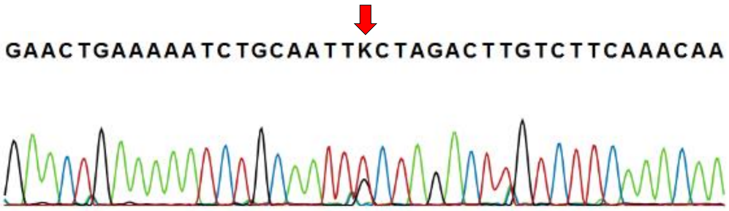


Cycle 40

2763 (T)

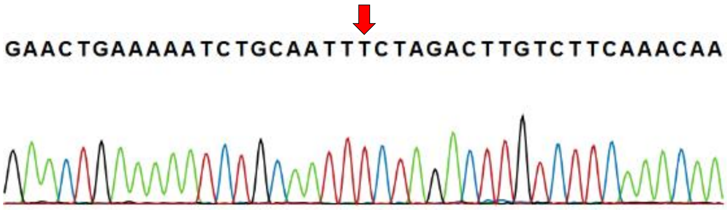

#### *ATH1* (339DelA) (Lys113fs)

**PCR: ATHP2for/ATHP2rev; sequencing: ATHP2for**

**Cycle 20**

**339 (A)**

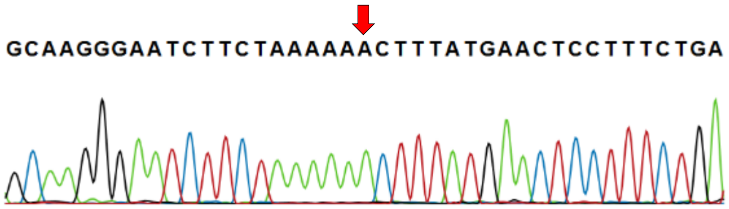


**Cycle 40**

**339 (A/C)**

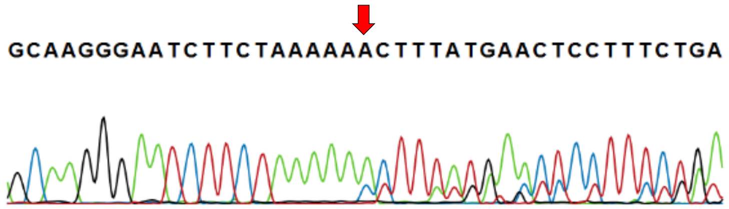


**Cycle 60**

**339 (A/C)**

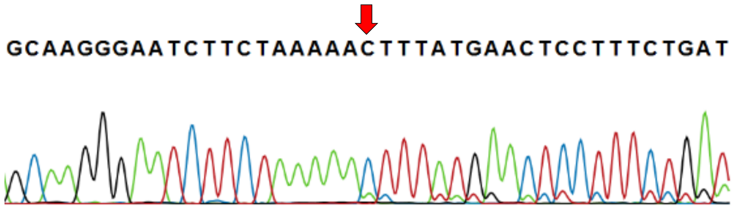


**Cycle 71 (end)**

**339 (C)**

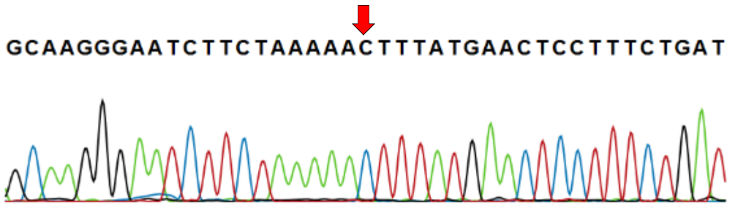


#### *RTT106* (650InsA) (Ile217fs)

**PCR: RTT106P2for/RTT106P2rev; sequencing: RTT106P2rev**

**Cycle 40**

**650 (T)**

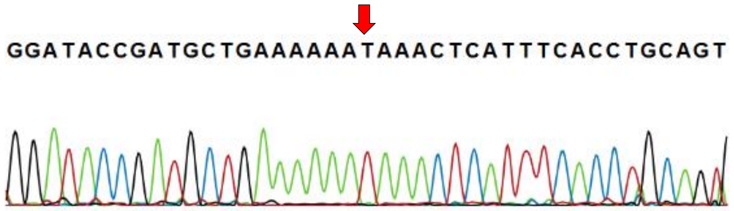


**Cycle 60**

**650 (A)**

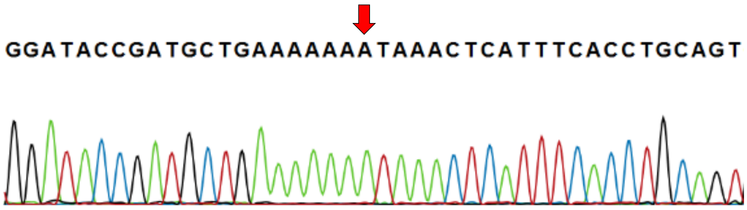

### <u>POPULATION 3 (P3)</u>

#### *ROM2* (G1318A) (Gly440Arg)

PCR: ROM2P3for/ROM2P3rev; sequencing: ROM2P3for

Progenitor
1318 (G)

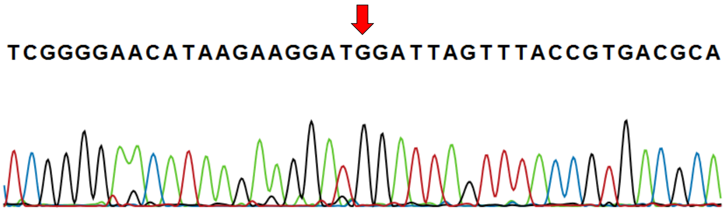


Cycle 20
1318 (A)

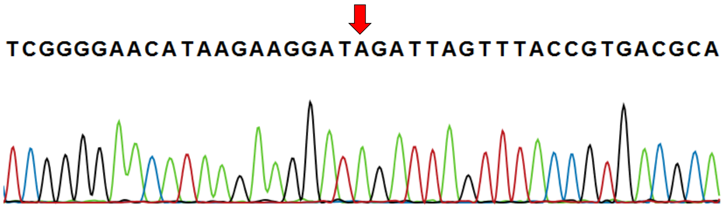


#### *MDS3* (G648T) (Arg216Ser)

PCR: MDS3P3for/MDS3P3rev; sequencing: MDS3P3for

Progenitor
648 (G)

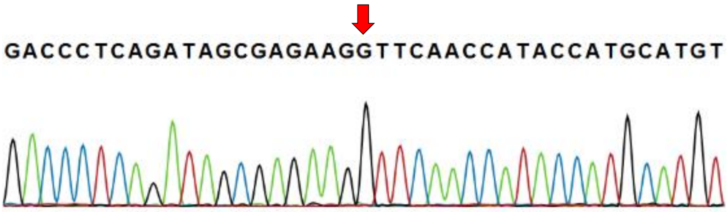


Cycle 20
648 (G/T)

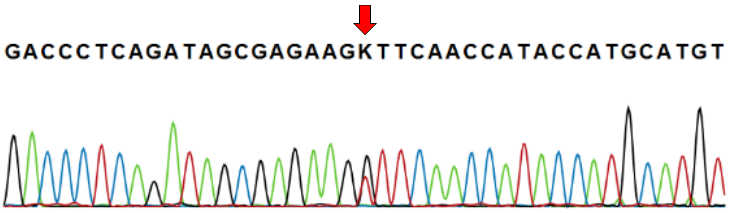


Cycle 40
648 (T)

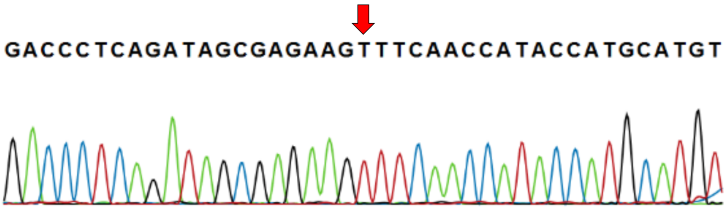

#### *NTH1* (117InsC) (Thr40fs)

**PCR: NTH1P3for/NTH1P3rev; sequencing: NTH1P3for**

**Cycle 20**

**117 (A)**

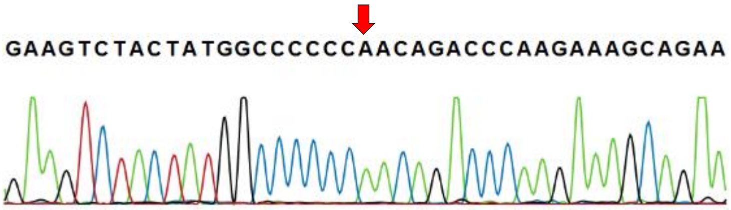


**Cycle 40**

**117 (C)**

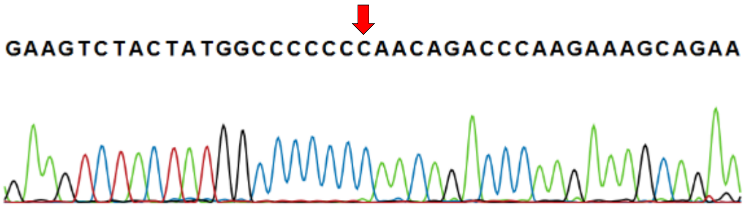


#### *ATH1* (G957A) (Trp319Stop)

**PCR: ATH1P3for/ATH1P3rev; sequencing: ATH1P3for**

**Cycle 20**

**957 (G)**

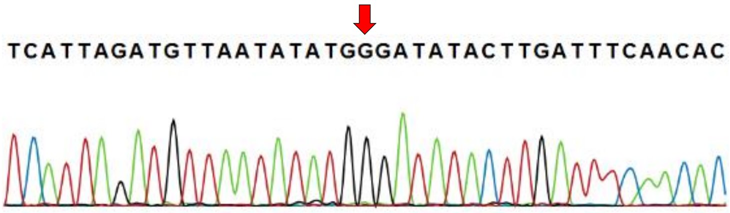


**Cycle 40**

**957 (A)**

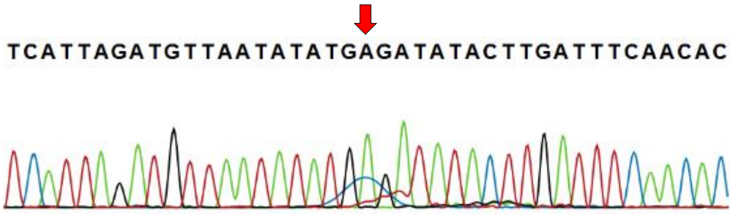


#### *USV1* (G182T) (Arg61Leu)

**PCR: USV1for/USV1rev; sequencing: USV1for**

**Cycle 60**

**182 (G)**

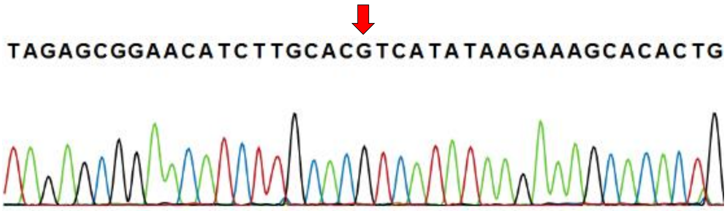


**Cycle 80**

**182 (T)**

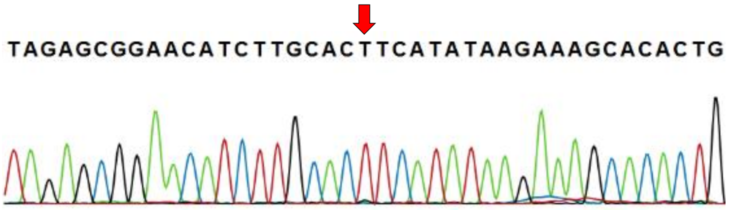

#### *PTR2* (C1436T) (Ser479Leu)

**PCR: PTR2for/PTR2rev; sequencing: PTR2rev**

**Cycle 40**

**1436 (C)**

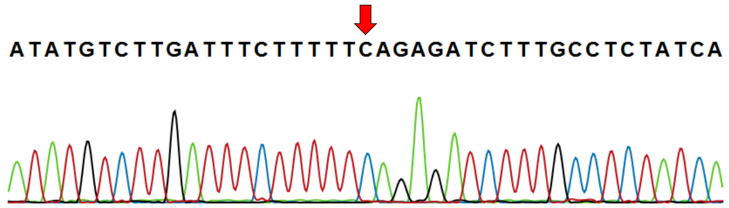


**Cycle 60**

**1436 (T/C)**

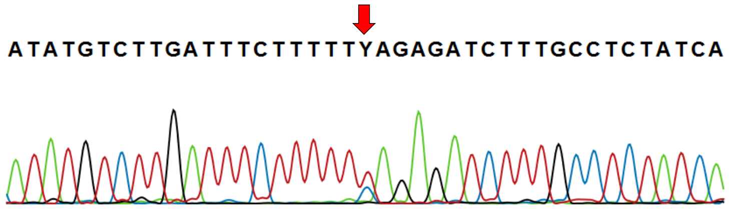


**Cycle 82 (end)**

**1436 (T)**

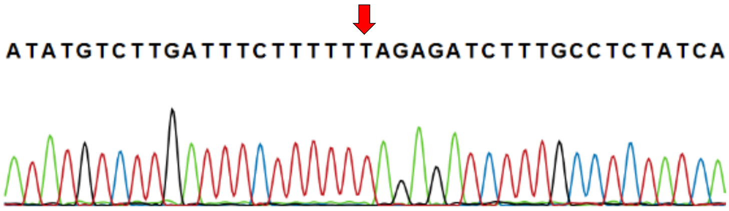


#### *PMD1* (C3039A) (Tyr1013Stop)

**PCR: PMD1P3for/PMD1P3rev; sequencing: PMD1P3for**

**Cycle 60**

**3039 (C)**

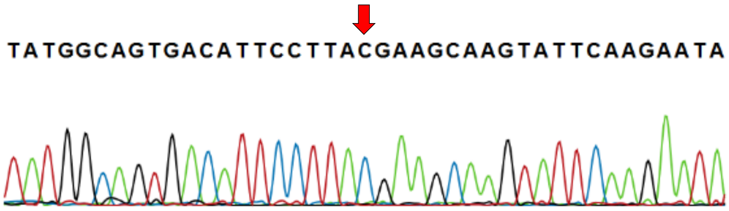


**Cycle 82 (end)**

**3039 (A)**

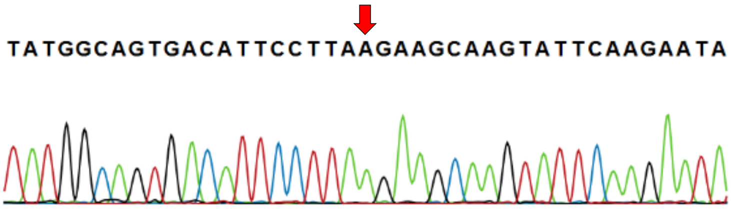


#### *IRA2* (5563DelT) (Cys1855fs), detected in P3c, but not in P3

**PCR: IRAP3for/RAP3rev; sequencing: IRAP3for**

**Progenitor**

**5563 (T)**

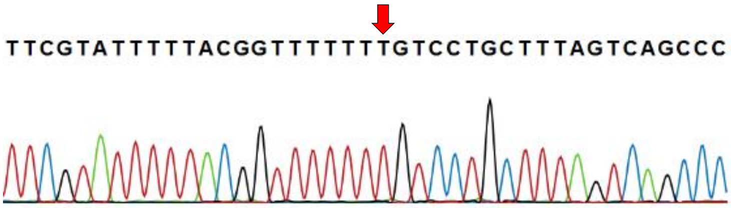


**Cycle 82 (end)**

**5563 (T)**

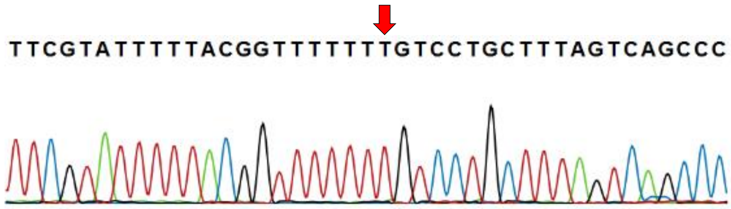

<u>**POPULATION 4 (P4)**</u>

#### *RAS2* (T2A) (Met1Lys)

**PCR: RASP4for/RASP4rev; sequencing: RASP4for**

**Progenitor**
**2 (T)**

AAAAAAAAAATGCCTTTGAACAAGTCGAACATAAGAGAGTA

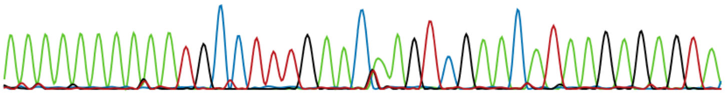

**Cycle 20**
**2 (A)**

AAAAAAAAAAAGCCTTTGAACAAGTCGAACATAAGAGAGTA

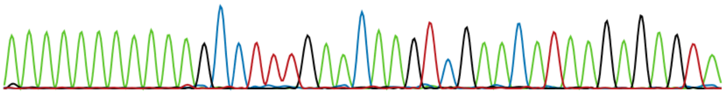

#### *PTR2* (T1084G) (Trp362Gly)

**PCR: PTR2for/PTR2rev; sequencing PTR2rev**

**Progenitor**
**1084 (T)**

TTATATTCTACCCAATTTATTGGACCCAATACGGTACCATG

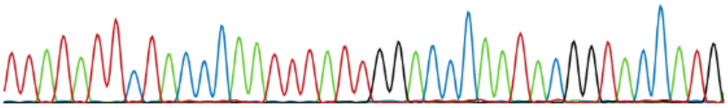

**Cycle 20**
**1084 (G)**

TTATATTCTACCCAATTTATGGGACCCAATACGGTACCATG

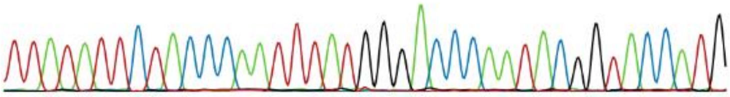

#### *BUD3* (2553DelT) (Phe851fs)

**PCR: BUD3P4for/BUD3P4rev; sequencing: BUD3P4for**

**Progenitor**
**2253 (T)**

AACAAAGGCAAAGGGTTTTTTGGCGTGTTAAAAAATGTTTT

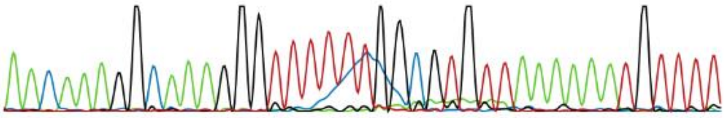

**Cycle 20**
**2253 (T/G)**

AACAAAGGCAAAGGGTTTTTTGGCGTGTTAAAAAATGTTTT

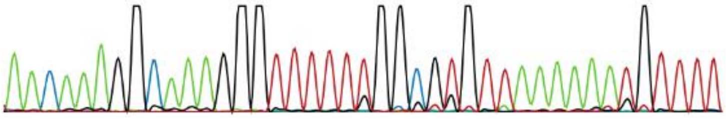

**Cycle 40**
**2253 (G)**

AACAAAGGCAAAGGGTTTTTGGCGTGTTAAAAAATGTTTTT

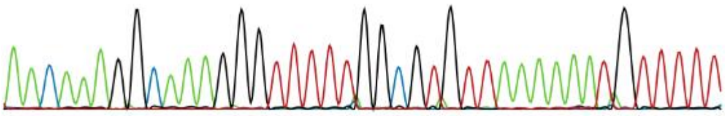

#### *APD1* (3’UTR +2 bp G>T)

PCR: APD1P4for/APD1P4rev; sequencing: APD1P4for

Cycle 40
3’UTR
+2 bp (G)

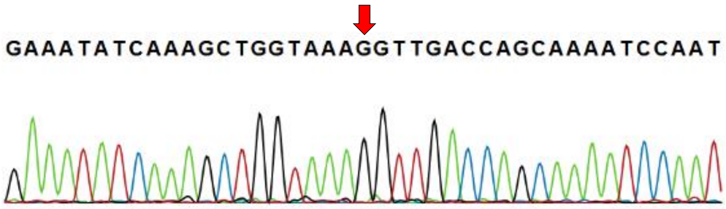


Cycle 60
3’UTR
+2 bp (T)

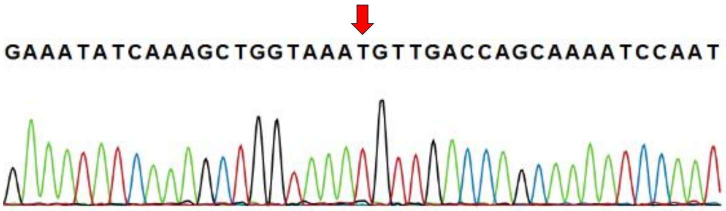


#### *Non-coding DNA between IPL1* and *SRP72* (Inversion: CCTTTGT> ACAAAGG)

PCR: IPL1for/IPL1re; sequencing: IPL1rev

Cycle 40
(CCTTTGT)

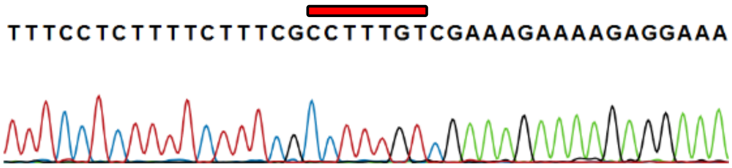


Cycle 60
(ACAAAGG)

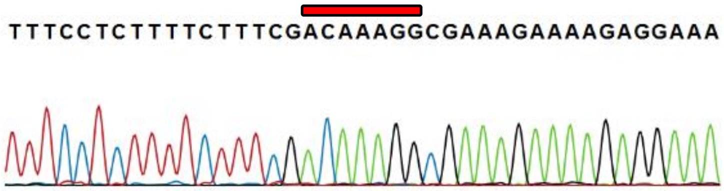
